# Novel mode of filament formation in UPA-promoted CARD8 and NLRP1 Inflammasomes

**DOI:** 10.1101/2020.06.27.175497

**Authors:** L. Robert Hollingsworth, Liron David, Yang Li, Andrew R. Griswold, Humayun Sharif, Pietro Fontana, Tian-Min Fu, Jianbin Ruan, Daniel A. Bachovchin, Hao Wu

## Abstract

NLRP1 and CARD8 are related cytosolic sensors that upon activation form supramolecular signalling complexes known as canonical inflammasomes, resulting in caspase-1 activation, cytokine maturation and/or pyroptotic cell death. NLRP1 and CARD8 use their C-terminal (CT) fragments containing a caspase recruitment domain (CARD) and the UPA subdomain of a function-to-find domain (FIIND) for self-oligomerization and recruitment of the inflammasome adaptor ASC and/or caspase-1. Here, we report cryo-EM structures of NLRP1-CT and CARD8-CT assemblies, in which the respective CARDs form central helical filaments that are promoted by oligomerized, but flexibly linked UPAs surrounding the filaments. We discover that subunits in the central NLRP1^CARD^ filament dimerize with additional exterior CARDs, which roughly doubles its thickness and is unique among all known CARD filaments. The thick NLRP1 filament only forms with the presence of UPA, which we hypothesize drives the intrinsic propensity for NLRP1^CARD^ dimerization. Structural analyses provide insights on the requirement of ASC for NLRP1-CT signalling and the contrasting direct recruitment of caspase-1 by CARD8-CT. Additionally, we present a low-resolution 4 ASC^CARD^–4 caspase-1^CARD^ octamer structure, illustrating that ASC uses opposing surfaces for NLRP1, versus caspase-1, recruitment. These structures capture the architecture and specificity of CARD inflammasome polymerization in NLRP1 and CARD8.

---

Innate immune pathways recognize and respond to a diverse array of intracellular threats. In one such pathway, cells identify and amplify danger signals through supramolecular signalling complexes called canonical inflammasomes^1,2^. Upon recognition of intracellular molecular patterns indicative of pathogens or endogenous damage, sensor proteins facilitate inflammasome assembly by undergoing large conformational changes that lead to their oligomerization^3,4^. Nucleotide binding domains (NBDs) often drive this self-oligomerization to cluster death-fold domains^5,6^, which in turn recruit downstream adapter and effector molecules. These death-fold domains, including pyrin domains (PYD) and caspase recruitment domains (CARD), participate in homotypic (CARD-CARD or PYD-PYD) interactions that ultimately lead to polymerization of caspase-1 into filaments^3,4,7–9^. This process increases the local concentration of caspase catalytic domains to facilitate its homo-dimerization and autoproteolysis, resulting in its activation^1–3^. Active caspase-1, a cysteine protease, then processes pro-inflammatory cytokines, including pro-IL-1β and pro-IL-18, to their bioactive forms and cleaves the pore-forming protein gasdermin-D (GSDMD) to promote cytokine release and often concomitant lytic cell death termed pyroptosis^10–12^.

Two unique sensor proteins, NLRP1 and CARD8, mediate inflammasome formation through their CARD-containing C-terminal fragment (CT) generated upon functional degradation of their respective N-terminal fragment (NT) by the proteasome^13,14^. This unusual mechanism of inflammasome activation is associated with autoproteolysis of the function-to-find domain (FIIND) common to NLRP1 and CARD8, which splits FIIND into ZU5 and UPA subdomains and results in noncovalently associated NT and CT^15,16^ (Fig. 1a). NLRP1-CT and CARD8-CT are therefore repressed by the NT until upstream cues induce degradation of the NT and release of the CT^17–20^ (Fig. 1b). Interestingly, one such cue is provided by small-molecule inhibitors of dipeptidyl peptidases DPP9 and DPP8, for which the mechanism of NLRP1 and CARD8 activation is unclear^21–23^. Activated CARD8 and NLRP1 inflammasomes are distinctive in that they are only composed of UPA and CARD, unlike others such as NLRC4 and NLRP3 inflammasomes which use NBDs and leucine-rich repeats (LRRs) to facilitate oligomerization and inflammasome activation^4–6,24^.

**Figure 1.**
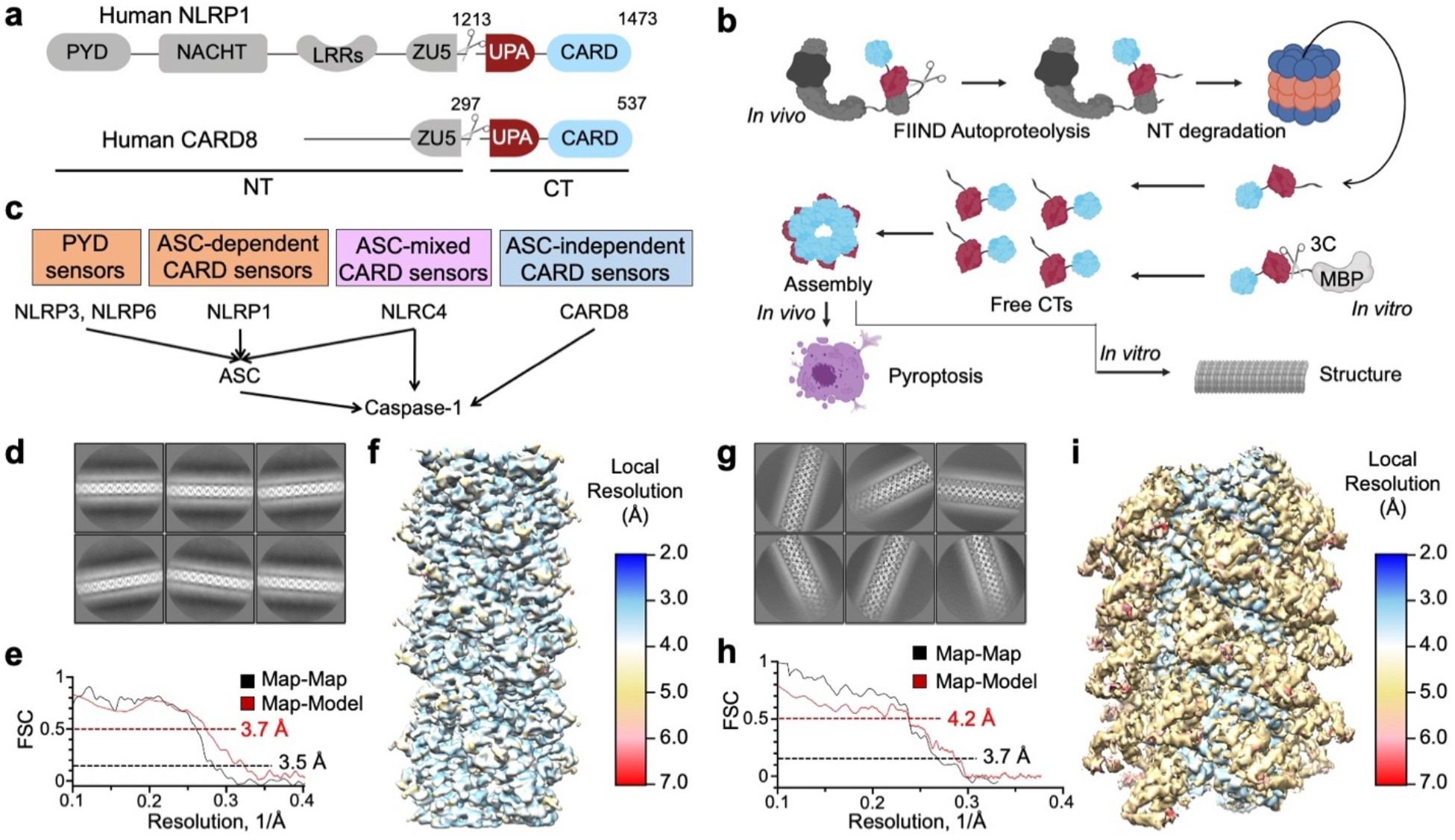
Structure determination of CARD8-CT and NLRP1-CT filaments. (**a**) Domain architecture of human NLRP1 and CARD8, each with a C-terminal domain (CT) containing UPA-CARD. (**b**) CARD8 and NLRP1 activation pathway in which signals leading to NLRP1 and CARD8 activation target them to the proteasome for N-terminal degradation. This leads to the release of free CT oligomers, which assemble the inflammasome. The purification strategy bypasses N-terminal degradation by directly expressing MBP-CT fusion proteins and cleaving the tag in vitro. (**c**) Classification of different inflammasome sensors by their signalling through the adaptor ASC. Human NLRP1 and CARD8 are ASC-dependent and ASC-independent, respectively. (**d**) Cryo-EM 2D classification of CARD8-CT filaments. (**e**) Gold-standard Fourier shell correlation (FSC) and map-model correlation plots of the CARD8-CT filament 3D reconstruction, which gave an overall resolution of 3.5 Å. (**f**) Local resolution of the CARD8-CT filament calculated by ResMap^61^. (**g**) Cryo-EM 2D classification of NLRP1-CT filaments. (**h**) FSC plots of the NLRP1-CT filament 3D reconstruction, which gave an overall resolution of 3.7 Å. (**i**) Local resolution of the NLRP1-CT filament calculated by ResMap^61^.

NLRP1 and CARD8 are highly expressed in a number of cell types and play important roles in both host defence and human diseases. Keratinocytes constitutively express high levels of NLRP1 and germline mutations of NLRP1 in humans lead to a number of skin-related inflammatory diseases, including multiple self-healing palmoplantar carcinoma (MSPC), familial keratosis lichenoides chronica (FKLC)^17^, vitiligo^25,26^, autoinflammation with arthritis and dyskeratosis (AIADK)^21,27^, and juvenile-onset recurrent respiratory papillomatosis (JRRP)^28^.

These mutations cause constitutive NLRP1 activation and downstream pyroptosis^17,21^, leading to damaging inflammation. For CARD8, the most highly expressing cell types are hematopoietic in origin, and DPP8/DPP9 inhibitor-induced pyroptosis via CARD8 is being pursued for treatment of acute myeloid leukemia^23^. Thus, understanding the molecular mechanisms governing NLRP1 and CARD8 inflammasome signalling will facilitate the discovery of new therapeutics for inflammatory diseases and cancers.

In addition to having the unique UPA subdomain, the specificity of NLRP1-CT and CARD8-CT for the CARD and PYD-containing adaptor ASC and for caspase-1 differ from most other inflammasome sensors (Fig. 1c). While ASC is a nearly universal adapter that bridges interactions between sensors and the CARD-containing caspase-1^1,29^, CARD8 does not interact with ASC and instead directly engages caspase-1^30^. In contrast to the CARD-containing protein NLRC4, which can activate caspase-1 both with and without ASC^6,31,32^, human NLRP1 is completely dependent on ASC and does not engage caspase-1 directly^30^.

The molecular bases for the assembly of UPA-CARD-mediated NLRP1-CT and CARD8-CT inflammasomes and for their differential specificity to ASC and caspase-1 are unknown. Here we determined the cryo-electron microscopy (cryo-EM) structures of NLRP1-CT and CARD8-CT filaments and analysed their assembly and specificity. Despite being invisible in the filament structures, the UPA domain is required for NLRP1 and CARD8 inflammasome signalling, suggesting that it promotes CARD clustering and filament formation and serves an analogous function to the NBD and LRRs in many other inflammasome sensors. The CARD filament of CARD8-CT resembles the helical filaments of caspase-1^CARD^, ASC^CARD^, and NLRC4^CARD^ ^7,33,34^ with certain structural variations. Surprisingly however, the CARD filament of NLRP1-CT is composed of CARD dimers in which an additional CARD flanks each subunit of the central CARD filament. Together with the low-resolution structure of an ASC^CARD^-caspase-1^CARD^ octamer, we discover new mechanisms of inflammasome formation and uncover the structural basis of hetero-oligomeric CARD-CARD interactions. During the preparation of this manuscript, a preprint for the structures of CARD8^CARD^ and NLRP1^CARD^ filaments was released^35^. While most conclusions of the study are similar to our study, the preprint did not report a CARD-CARD dimer in the NLRP1^CARD^ filament structure. We speculate that subtle differences in constructs and sample preparation led to the observed structural differences, and together our studies bolster the conclusion that the UPA subdomain on the NLRP1 and CARD8 CTs facilitates inflammasome assembly and signalling.

## Results

### Cryo-EM structure determination of CARD8 and NLRP1 CARD filaments

In general, inflammasomes leverage supramolecular filamentous structures to nucleate the polymerization of caspase-1, which in turn increases its local concentration to facilitate activation^3,7,33^. To elucidate if and how CARD8-CT and NLRP1-CT form filaments, we expressed these CTs in fusion with an N-terminal maltose-binding protein (MBP) tag separated by a linker cleavable by the human rhinovirus (HRV) 3C protease. We posited that such a bulky tag would disrupt oligomerization and facilitate purification of monomeric proteins for controlled CT filament formation *in vitro* (Fig. 1b, Supplementary Fig. 1-2). However, the MBP-fusion protein still formed small oligomers. By systematically optimizing cleavage conditions, we purified short (~100-200 nm) filaments that still contained some uncleaved MBP-tagged proteins. Despite incorporation of MBP-tagged subunits, these small filaments behaved well on cryo-EM grids and enabled the calculation of CARD8-CT and NLRP1-CT maps at resolutions of 3.5 Å and 3.7 Å respectively (Fig. 1d-i, Supplementary Fig. 1-2, Supplementary Table 1). Unexpectedly, 2D classifications and 3D reconstructions revealed both similarities and differences between these two filament structures (see below).

### Structure of the CARD8-CT filament

Despite being included in the construct, no UPA density was observed in the CARD8-CT filament structure, in which only the CARD8^CARD^ filament was visible (Fig. 2a-b). One possible explanation is that CARD8^UPA^ does not follow the CARD helical symmetry but is orderly associated with the central CARD8^CARD^ filament. However, subtracting out the central density followed by 3D classification without symmetry did not reveal any new density. Given the predicted 17-residue unstructured linker between UPA and CARD, we proposed that CARD8^UPA^ and any residual MBP molecules must be flexibly linked to the core CARD8^CARD^ filament and were averaged out during data processing.

**Figure 2.**
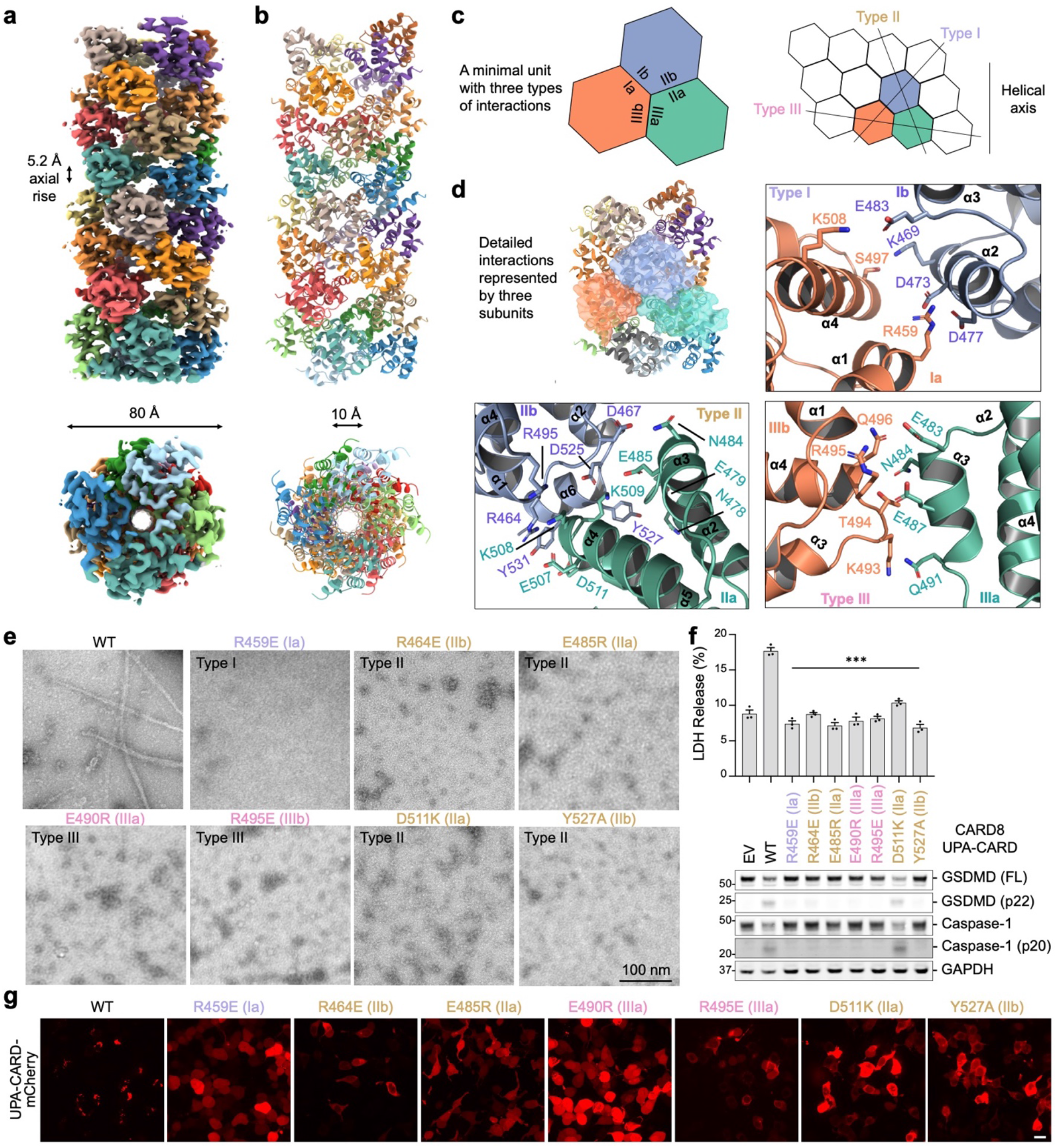
Cryo-EM structure and mutagenesis of the CARD8-CT filament. (**a-b**) Overall structure of the CARD8-CT inflammasome, with a resolved CARD8^CARD^ filament. The cryo-EM density (**a**) and atomic model (**b**) are colored by subunit. (**c**) Illustration of classical CARD filament interfaces: type I, II, and III. (**d**) Detailed CARD−CARD interactions in the CARD8-CT filament structure colored as in (**c**). (**e**) Negative stain electron microscopy snapshots of CARD8^CARD^ mutants that abolish filament formation *in vitro*. (**f**) Inflammasome assay of CARD8-CT-mCherry mutants transiently expressed in HEK293T cells that stably express caspase-1 and GSDMD. Filament-deficient mutants abolish inflammasome activity as measured by LDH release (top) and western blot (bottom) for activated inflammasome components. *** p < 0.001 compared to WT by two-sided Student’s t-test. Data are means ± SEM of three biological replicates. (**g**) Confocal imaging of HEK293T cells transiently expressing CARD8^UPA-CARD^-mCherry WT and mutants. Filament-deficient mutants abolish filament formation in cells. Experiments were performed with two biological replicates. Scale bar 5μM. In (**e-g**), mutations are labeled in the colors of the interface types as defined in (**c**).

Like other CARD filament structures^4,7,33,34,36,37^, the CARD8^CARD^ filament also possesses a one-start helical symmetry, with a refined left-handed rotation of 99.1° and an axial rise of 5.2 Å per subunit (~3.6 subunits per helical turn). The filament has a dimeter of ~80 Å with a central hole of ~10 Å (Fig. 2a-b). Analysis of the three types of asymmetric interactions (Fig. 2c) that are characteristic of death-fold filaments^38^ revealed that the type I interaction is unusually small, with only ~150 Å^2^ buried surface area per partner, in comparison to ~350 Å^2^ and ~560 Å^2^ for those in ASC^CARD^ and caspase-1^CARD^ filaments. In contrast, the type II interface is substantially more extensive, burying ~490 Å^2^ surface area per partner, and the type III interface covers ~230 Å^2^ per partner. Because of the small type I interface, the filament structure looks almost perforated with visible gaps (Fig. 2a).

Detailed inspection of the three interfaces revealed many charge-charge pairs as well as other hydrophilic interactions including R549 at type Ia with D473 and D477 at type Ib, K509 of type IIa with D525 of type IIb, E507 of type IIa with R464 of type IIb, N478 and E479 of type IIa with Y527 of type IIb, and E483 and E487 of type IIIa with R495 of IIIb (Fig. 2d). We leveraged the structure to design point mutations that would abolish CARD filament formation. We used a filament formation assay, in which we cleaved recombinant MBP-tagged CARD8^CARD^ protein overnight, followed by EM imaging of the negatively stained sample. While the wild-type protein formed filaments when the bulky MBP tag was removed, seven different point mutants, one at the type I interface, four at the type II interface and two at the type III interface, completely abolished filament formation *in vitro* (Fig. 2e). Of note, unlike many CARDs that form filaments at low μM concentrations^7,33^, we had to raise the concentration of CARD8^CARD^ to 15 μM to see consistent filaments in multiple grid areas under negative stain EM (Supplementary Fig. 3).

Next, we employed a reconstituted HEK293T cell system stably expressing caspase-1 and GSDMD to assess whether these mutants were able to signal in cells^23^. We transfected these cells with plasmid encoding for WT or mutant CARD8-CT (UPA-CARD), and 24 h later, analysed the supernatant for lactate dehydrogenase (LDH) activity, a hallmark of pyroptotic cell death, and the lysate by immunoblotting for caspase-1 and GSDMD cleavage (Fig. 2f). As expected, WT CARD8-CT induced elevated LDH activity, and exhibited prominent caspase-1 and GSDMD cleavage. In contrast, six out of the seven filament-deficient mutants showed background level LDH release with no discernible caspase-1 or GSDMD cleavage, similar to the empty vector (EV). Only the D511K mutant displayed caspase-1 and GSDMD cleavage but still significantly impaired LDH release; this discrepancy suggested that immunoblotting may be less quantitative and that LDH release is more indicative of the mutational effects. Furthermore, we expressed UPA-CARD as a mCherry fusion protein in HEK293T cells and found that although WT formed strong punctate structures indicative of filament formation, all the UPA-CARD constructs with CARD interface mutations showed diffuse distribution consistent with defective filament formation (Fig. 2g). Thus, these cellular data confirmed that CARD-CARD interactions in the filament are crucial for UPA-CARD-mediated inflammasome signalling.

### Specificity of the CARD8-CT filament for caspase-1

While most CARD-containing inflammasome proteins, including human NLRP1 and NLRC4, amplify signalling through the adapter protein ASC^30,32^, CARD8 is the only known inflammasome sensor that cannot engage ASC and instead exclusively binds caspase-1^30^. To address this specificity, we first investigated the surface charges of CARD8, ASC, and caspase-1 filaments in hypothetical complexes. In the orientation of the CARD8^CARD^ filament shown (Fig. 2b), if ASC^CARD^ or caspase-1^CARD^ filament is placed above the CARD8^CARD^ filament layer, the surface charges at the interfaces would have been all largely negative and these surfaces should repel each other (Fig. 3a). When the caspase-1^CARD^ filament is placed below the CARD8^CARD^ filament layer, the charge complementarity between CARD8^CARD^ and caspase-1^CARD^ is apparent, especially with strong positive (caspase-1) and negative (CARD8) patches both mainly locating at the outer side of the filament cross-sections (Fig. 3b). In contrast, the same placement for a hypothetical CARD8^CARD^−ASC^CARD^ complex reveals that the positive charge in ASC is near the centre of the filament, which does not complement the peripherally localized negative charge on the CARD8 filament (Fig. 3b).

**Figure 3.**
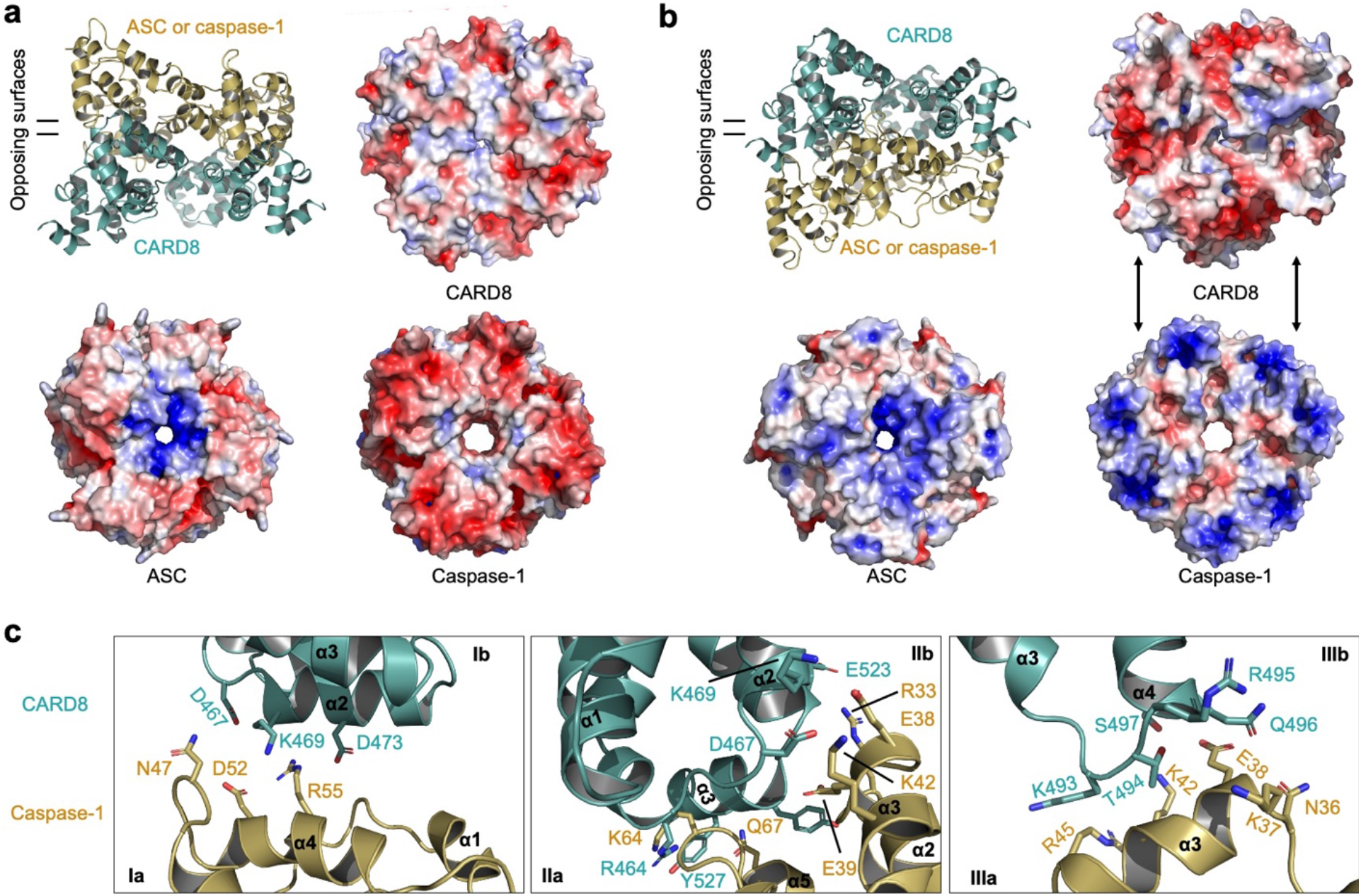
Specificity of the CARD8-CT filament for caspase-1. (**a**) A modeled octamer between a layer of ASC or caspase-1 (gold) on top of a CARD8 layer (green). Electrostatic surfaces on the interaction interface are shown, indicating that negatively charged surfaces for ASC and caspase-1 likely do not match the negatively charged CARD8 surface. (**b**) A modeled octamer between a layer of ASC or caspase-1 (gold) below a CARD8 layer (green). Electrostatic surfaces on the interaction interface are shown, indicating that CARD8 is likely compatible with caspase-1 given the complimentary surface charge, but not with ASC. This explains why CARD8 cannot recruit ASC and has specificity for caspase-1. (**c**) Detailed modelled CARD-CARD type I-III interactions between CARD8 (green) and caspase-1 (gold). All electrostatic surfaces were generated with +/− 5 kT/e.

To further elucidate the structural basis for the CARD8-caspase-1 interaction, we analysed the modelled interfaces type by type (Fig. 3c). The modelled CARD8−caspase-1 interfaces are extensive, with calculated buried surface areas of ~150, 600, 250 Å^2^ per partner for type I, II and III interfaces respectively. Consistent with the surface charge analysis, the interfaces are dominated by charged pairs including K469 of CARD8 to D52 of caspase-1 and D473 of CARD8 to R55 of caspase-1 at the type I interface, E523 of CARD8 to R33 of caspase-1, D467 and K469 of CARD8 to E38 and K42 of caspase-1, and a cluster of interactions involving Y527 and Y531 of CARD8 and K64 and Q67 of caspase-1 at the type II interface, and K493-S497 of CARD8 to K37, E38, K42 and R45 of caspase-1 at the type III interface (Fig. 3c). Thus, structural analysis confirmed favourable interactions between CARD8 and caspase-1.

### Structure of the NLRP1-CT filament containing CARD dimers

Similar to the CARD8-CT filament structure, no UPA density was observed in the NLRP1-CT filament structure, suggesting that the UPA subdomain, with the 25-residue predicted unstructured linker to the CARD, is also flexibly connected to the core NLRP1^CARD^ filament (Fig. 4a). The NLRP1^CARD^ filament has one-start helical symmetry, with a left-handed rotation of 100.8° and an axial rise of 5.1 Å per subunit (~3.6 subunits per helical turn), similar to other CARD filament structures^4,7,33,34,36,37^. Strikingly however, unlike any other CARD filament structures known to date, the NLRP1^CARD^ filament is composed of NLRP1^CARD^ dimers, rather than monomers (Fig. 4a-b), which is also reflected in the thicker dimensions of NLRP1-CT filaments than CARD8-CT filaments in both 2D classes and the 3D volume (Fig. 1d, 1f, 1g, 1i). The inner NLRP1^CARD^ filament is equivalent to other CARD filament structures, and for all NLRP1^CARD^ subunits in the inner filament, the dimerically related NLRP1^CARD^ subunits form the outer layer of the NLRP1^CARD^ filament (Fig. 4a). The total diameter of the NLRP1^CARD^ filament is approximately 140 Å, with an inner hole of ~10 Å (Fig. 4a).

**Figure 4.**
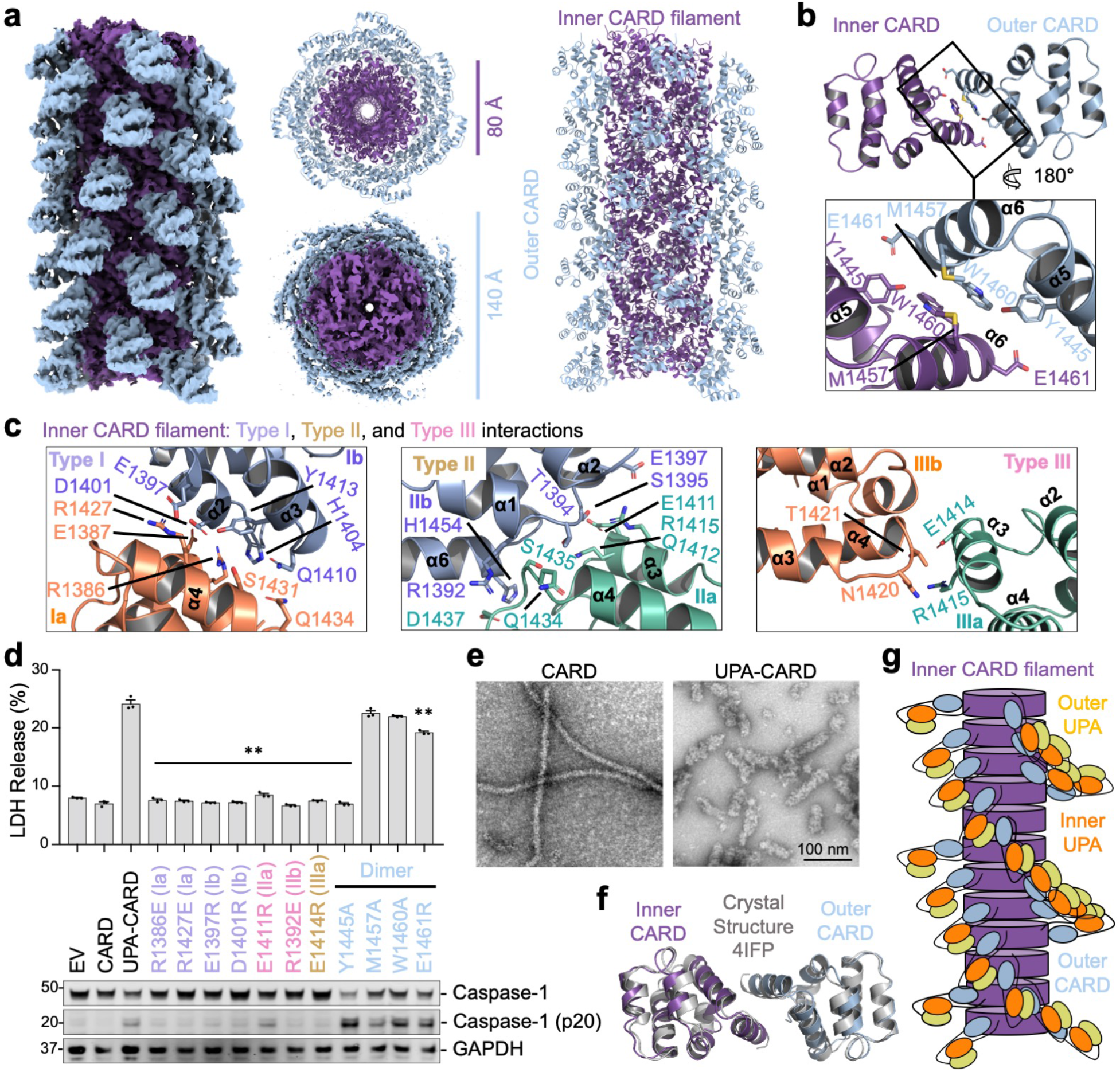
Cryo-EM structure of the dimeric NLRP1 CARD filament. (**a**) Overall structure of the NLRP1-CT inflammasome. Each subunit of the inner helical CARD filament (purple) dimerizes with an outer CARD (blue). (**b**) Dimeric interface between an inner and outer CARD pair. (**c**) Zoom-ins of type I, II, and III helical interfaces between inner CARD molecules. Colored as in Fig. 2c. (**d**) Structure-guided mutations of helical interfaces and the CARD-CARD dimer interface on UPA-CARD inflammasome signalling. LDH release (top) and western blot (bottom) are shown. ** p < 0.01 compared to UPA-CARD by two-sided Student’s t-test. Data are means ± SEM of three biological replicates. (**e**) EM of negatively stained NLRP1^CARD^ filaments with a diameter of ~10 nm (left) and thicker NLRP1^UPA-CARD^ filaments with a diameter of ~20 nm (right). Scale bar 100 nm. (**f**) Alignment between Inner-outer CARD (purple and blue) and a dimer within the crystallographic symmetry unit of the NLRP1 CARD crystal structure (grey). (**g**) Proposed model for NLRP1^UPA-CARD^ filament formation. A central CARD filament is decorated by outer CARD subunits, with adjacent UPAs (not visible in the structure) promoting oligomerization.

The NLRP1^CARD^ dimer is mediated by reciprocal interactions at helices α5 and α6, burying a substantial ~300 Å^2^ surface area per partner, and involving residues with large side chains such as Y1445, M1457, W1460, and E1461 (Fig. 4b). These regions of helices α5 and α6 are not involved in the inner core filament interaction, which instead is mediated by the classical type I, II, and III CARD-CARD interactions (Fig. 4c). Different from the CARD8^CARD^ filament, the type I interface in the NLRP1^CARD^ filament is more extensive, with type I, II and III interfaces burying ~300, 430 and 140 Å^2^ surface area per partner, respectively. Charged and hydrophilic interactions dominate the interactions, including R1386 of type Ia with D1401, Y1413 and H1404 of type Ib, R1427 of type Ia with E1397 and D1401 of type Ib, E1411 of type IIa with S1395 of type IIb, Q1434 and D1437 of type IIa with R1392 of type IIb, and E1414 of type IIIa with T1421 of type IIIb (Fig. 4c).

To elucidate the role of the observed NLRP1^CARD^ filament in signalling, we employed the same reconstituted HEK293T cell system stably expressing caspase-1 and GSDMD used for assessing CARD8 mutants in cells (Fig. 2f). We transfected these cells with plasmids encoding WT or mutant NLRP1 UPA-CARD in addition to ASC, and 24 h later, analysed the supernatant for LDH activity, and the lysate for caspase-1 cleavage (Fig. 4d). In contrast to WT UPA-CARD, structure-designed mutants including R1386E (Ia), R1427E (Ia), E1397R (Ib), D1401R (Ib), R1392E (IIb), and E1414R (IIIa) compromised cell death measured by both LDH release and caspase-1 processing (Fig. 4d). The E1411R (IIa) mutant displayed a certain level of caspase-1 processing but was almost completely defective in LDH release (Fig. 4d). Together, the mutational analysis demonstrated the validity of the filament structure in NLRP1 signalling.

### NLRP1 UPA promotes CARD dimerization, CARD helical oligomerization and signalling

To elucidate the function of the NLRP1 UPA subdomain as well as CARD dimerization, we compared the signalling activity of WT CARD, WT UPA-CARD, and UPA-CARD with several CARD dimerization mutants (Fig. 4d). To the contrary of WT UPA-CARD, the CARD alone construct showed background level LDH release and no caspase-1 processing, similar to the empty vector (EV). Among the UPA-CARD constructs with CARD dimerization mutations, Y1445A compromised LDH release but retained caspase-1 processing, suggesting partial defectiveness. However, several other dimerization mutants, M1457A, W1460A, and E1461R, showed no discernible impact on inflammasome signalling. Thus, while the UPA is required for UPA-CARD inflammasome signalling, the dimer likely promotes assembly to a lesser degree. This result is consistent with lack of strict sequence conservation of these residues among NLRP1 from different species, and with Y1445 as the most conserved residue at the dimerization interface (Supplementary Fig. 4).

The functional role of UPA in inflammasome signalling is suggestive of its ability to dimerize or oligomerize such that its presence promotes CARD filament formation. This ability of UPA is supported by negative staining EM analysis of CARD filaments versus UPA-CARD filaments, both formed *in vitro* using recombinant protein. While UPA-CARD formed thick filaments, CARD alone only formed thin filaments that are characteristic of the inner CARD (Fig. 4e) and of other published CARD filaments^4,7,33,34,36,37^. These data suggest that UPA dimerization or oligomerization promotes NLRP1 CARD dimerization, as well as NLRP1 helical filament formation, strengthening the role of UPA in the NLRP1 signalling paradigm.

If UPA mediates NLRP1 CARD dimerization, why does the CARD8-CT fail to induce CARD8^CARD^ dimerization? To address this question, we inspected the crystal packing interactions in the previously determined MBP-fused NLRP1^CARD^ structure (PDB ID: 4IFP) and MBP-fused CARD8^CARD^ structure (PDB ID: 4IKM) to see if a dimer was observed^39,40^. The MBP fusion kept NLRP1^CARD^ and CARD8^CARD^ from forming filaments. For NLRP1, all three independent molecules in the crystallographic asymmetric unit form a symmetrical dimer in the crystal lattice, which superimposes well with the dimer observed in the filament (Fig. 4f). These analyses suggest that NLRP1^CARD^ has an intrinsic propensity to form dimers. In contrast, no CARD8^CARD^ dimer was observed in crystal lattice. Further, the dimerization interface in NLRP1 is not conserved in CARD8 (Supplementary Fig. 4). Based on these data and analyses, we propose a model of UPA-induced NLRP1^CARD^ filament formation in which flexibly linked UPA dimers or oligomers promote the intrinsic tendency of NLRP1^CARD^ dimerization and its helical polymerization to mediate inflammasome formation (Fig. 4g).

### Specificity of the NLRP1-CT filament for ASC

In contrast to the specificity of CARD8 for caspase-1, NLRP1-CT only engages ASC, but not caspase-1 ^30^. To address this specificity, we first investigated the surface charges of NLRP1, ASC, and caspase-1 filaments in hypothetical complexes. In the side view orientation of the NLRP1^CARD^ filament shown (Fig. 4a), we tested two alternative models in which the ASC^CARD^ or caspase-1^CARD^ filament is placed either above (Fig. 5a) or below the NLRP1^CARD^ filament layer (Fig. 5b). Among these models, the best match by visual inspection of the charge complementarity of the cross sections is between NLRP1 and ASC when NLRP1^CARD^ is placed above ASC^CARD^, with centrally localised negative charge on NLRP1^CARD^ and centrally localised opposing positive charge on ASC^CARD^ (Fig. 5b).

**Figure 5.**
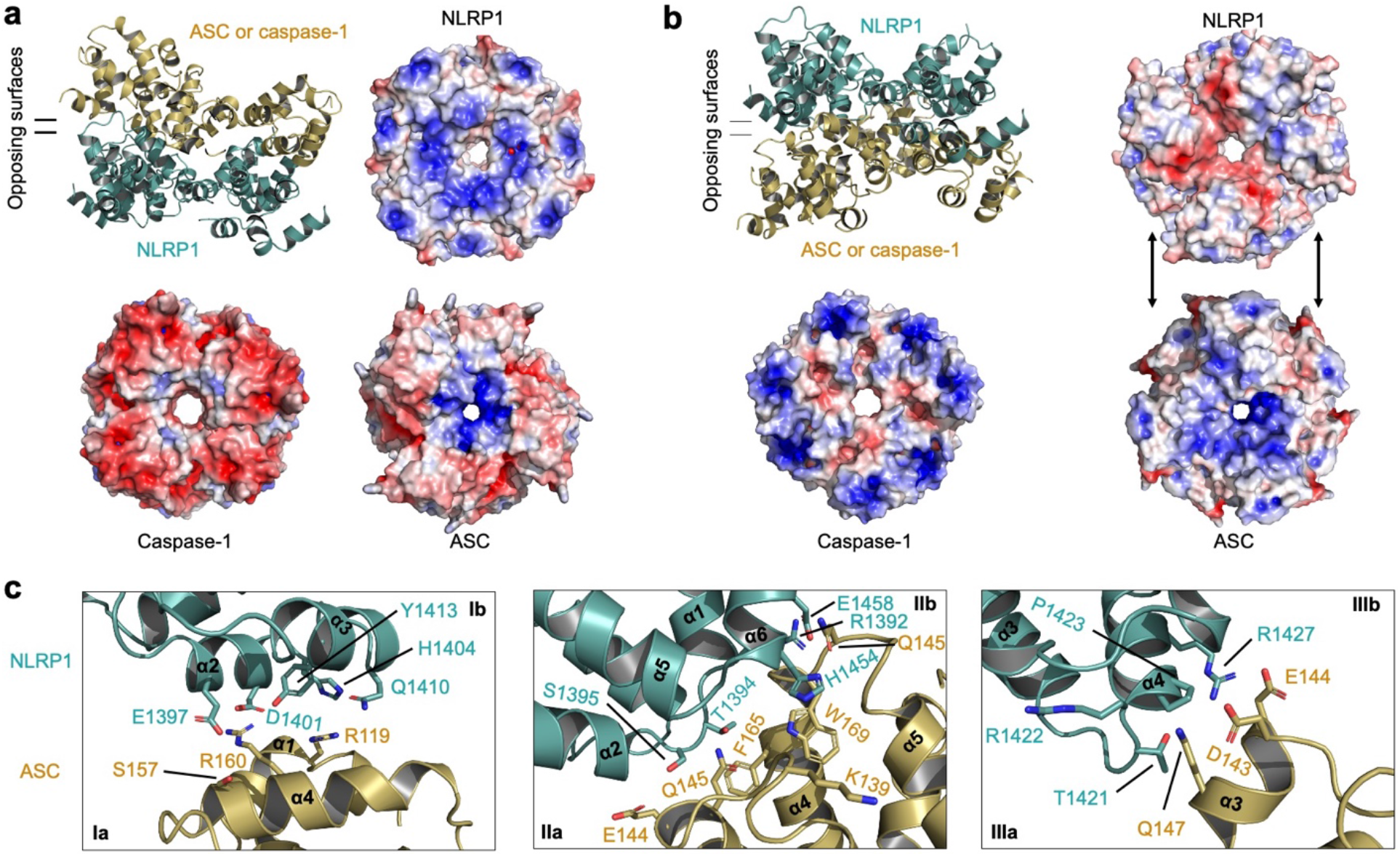
Specificity of the NLRP1-CT filament for ASC. (**a**) A modelled octamer between a layer of ASC or caspase-1 (gold) on top of an NLRP1 layer (green). Electrostatic surfaces on the interaction interface are shown. NLRP1 and ASC surfaces both have positive changes near the centre and should be incompatible. Caspase-1 surface has negative charges throughout whereas NLRP1 surface does not appear to have enough counter positive charges to be compatible. (**b**) A modelled octamer between a layer of ASC or caspase-1 (gold) below an NLRP1 layer (green). Electrostatic surfaces on the interaction interface are shown, indicating that NLRP1, with its negatively charges near the centre, is likely compatible with ASC, with its positive changes also near the centre. There is no such charge complementarity between NLRP1 and caspase-1, explaining why NLRP1 cannot recruit caspase-1 and has specificity for ASC. (**c**) Detailed modelled CARD-CARD type I-III interactions between NLRP1 (green) and ASC (gold). All electrostatic surfaces were generated with +/− 5 kT/e.

To further elucidate the structural basis for the NLRP1−ASC interaction, we analysed the modelled interfaces type by type. The modelled NLRP1−ASC interfaces are extensive, with calculated buried surface area of ~170, 500, 190 Å^2^ per partner for type I, II and III interfaces respectively. Consistent with the surface charge analysis, the interfaces are dominated by charged pairs and hydrophilic pairs including E1397 and D1401 of NLRP1 with R119 and R160 of ASC at the type I interface, T1394 and S1395 of NLRP1 with E144 and Q145 of ASC at the type II interface, and R1427 of NLRP1 with D143 and E144 of ASC at the type III interface (Fig. 5c). Thus, the structural analysis confirmed favourable interactions between NLRP1 and ASC.

### ASC filament uses a different surface for caspase-1 recruitment

We predicted previously that ASC^CARD^ uses its type Ib, IIb, and IIIb surfaces to interact with the caspase-1^CARD^ type Ia, IIa, and IIIa surfaces, respectively^33^. To test this prediction, we designed the type IIa W169G mutant of ASC^CARD^, and the type IIb G20K mutant of caspase-1^CARD^ based on the ASC and caspase-1 filament structures^7,33^ and connected them in one polypeptide chain using a 5 x GSS linker (Fig. 6a). As expected, these mutations did not impair the formation of an ASC^CARD^−caspase-1^CARD^ oligomer (Supplementary Fig. 5a), and multi-angle light scattering (MALS) measurement revealed a main peak of molecular mass of 86.3 kDa, corresponding to a tetramer of the ASC^CARD^−caspase-1^CARD^ fusion protein, or an octamer of CARDs with 4 ASC^CARD^ molecules and 4 caspase-1^CARD^ molecules (Fig. 6b).

**Figure 6.**
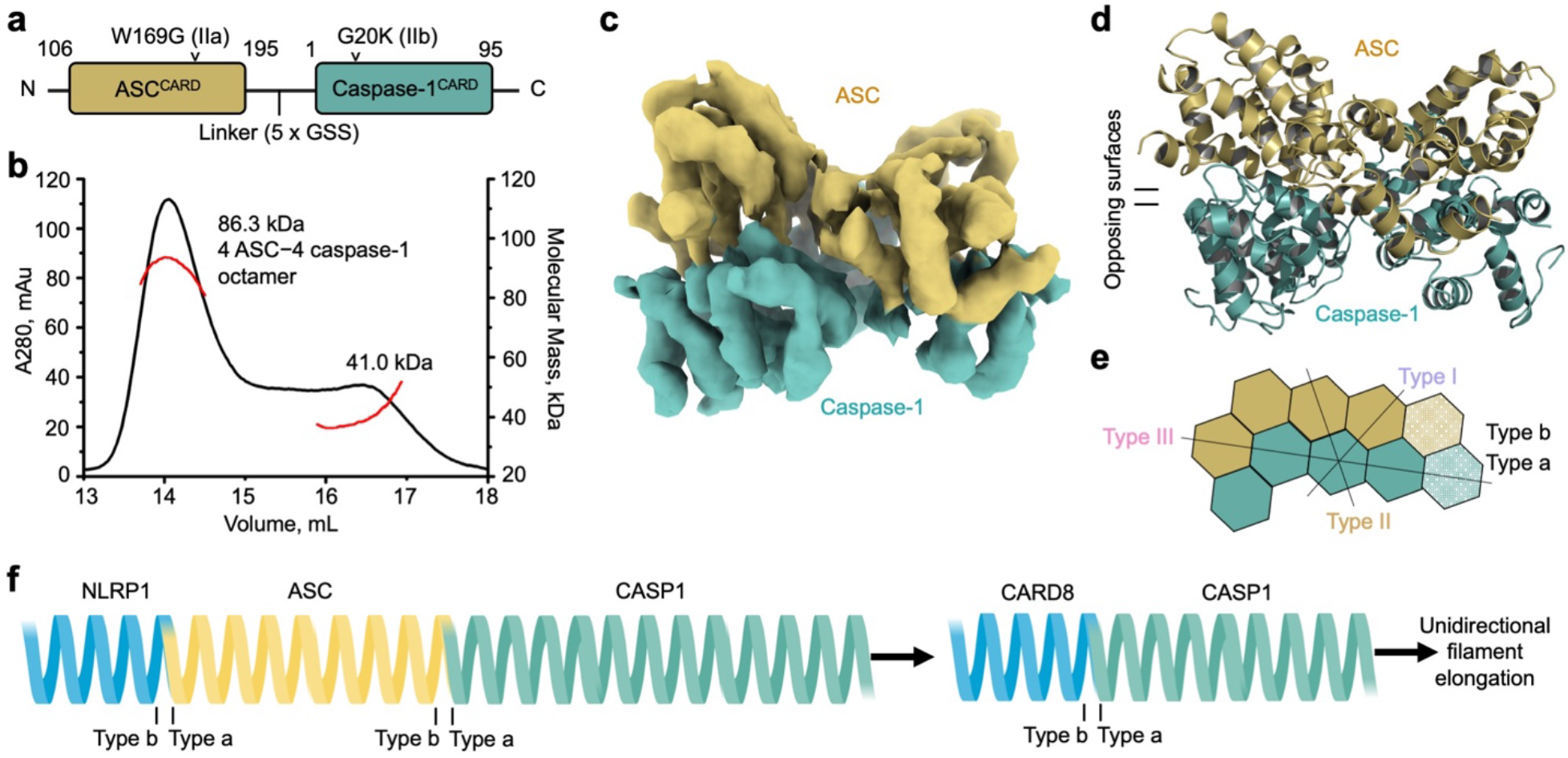
Purification and structure determination of an ASC^CARD^−caspase-1^CARD^ octamer. (**a**) Construct design of ASC^CARD^ and caspase-1^CARD^ linked by 5 x GSS linker. (**b**) MALS data for the ASC^CARD^−caspase-1^CARD^ complex, with a molecular mass of 86.3 kDa corresponding to a complex formed by 4 subunits of ASC^CARD^ and 4 subunits of caspase-1^CARD^. (**c**) Cryo-EM density of the ASC^CARD^−caspase-1^CARD^ octamer at 5.0 Å resolution in which a layer of ASC^CARD^ (gold) is located on top of a caspase-1^CARD^ layer (green). (**d**) Cryo-EM model of the ASC^CARD^−caspase-1^CARD^ octamer. (**e**) Schematic diagram of the octamer complex with accurately depicted interfaces. (**f**) Simplified illustration of NLRP1 and CARD8 hierarchy in inflammasome signalling. NLRP1 recruits ASC, followed by capase-1 recruitment. In contrast, CARD8 can only recruit caspase-1 directly. Both signalling pathways rely on unidirectional filament polymerization.

Despite the relatively small size, we collected a cryo-EM dataset and found that the 2D classes were quite detailed (Supplementary Fig. 5b-d). We then determined the 3D structure of the ASC−caspase-1 octamer at a resolution of 5.0 Å calculated by FSC between half maps (Supplementary Fig. 5e). The eight CARD structures are apparent in the map, and the ASC^CARD^ and caspase-1^CARD^ structures are distinguishable by the shorter α6 in caspase-1^CARD^, allowing unambiguous placement of ASC^CARD^ and capase-1^CARD^ subunits into the density (Fig. 6c-d). In this assembly, there is one type I interaction and three type III interactions within ASC subunits and within caspase-1 subunits, and there are three type I interactions, four type II interactions and one type III interaction between ASC and caspase-1 (Fig. 6e), which must have represented the optimal mode of interaction between ASC and caspase-1.

In all three types of interactions between ASC and caspase-1, ASC always uses the type b surfaces (Ib, IIb, and IIIb) to interact with the type a surfaces (Ia, IIa and IIIa) of caspase-1 (Fig. 6e). By contrast, ASC uses its type a surfaces for recruitment by NLRP1, suggesting a hieratical inflammasome formation that proceeds through an ordered manner from NLRP1, to ASC, then to caspase-1, with potential signal amplification (Fig. 6f). This ordered assembly is made possible by the use of two opposing ASC filament surfaces with specificity towards either NLRP1 or caspase-1. However, given the oligomerization ability of the PYD of ASC, it is also possible that the same ASC molecule does not necessarily need to interact with both NLRP1 and caspase-1. In this case, the NLRP1^CARD^-ASC^CARD^ oligomer facilitates ASC^PYD^ filament assembly, and peripheral ASC^CARD^ facilitates bundling into a single perinuclear punctum and caspase-1^CARD^ recruitment, like for NLRC4^41^. In either scenario, these data indicate that ASC uses opposing surfaces for interaction with NLRP1 and caspase-1.

## Discussion

The death-fold domain superfamily is widely represented in innate immune signalling pathways and mediates homo- and hetero-oligomerization through filament formation^4,42,43^. In this study, we showed that NLRP1-CT and CARD8-CT also use their CARD to assemble filamentous structures for inflammasome assembly. We posit that these CARD filaments are decorated by the flexibly linked UPA subdomains (Fig. 4g), and that UPA oligomerization decreases the threshold for filament assembly by locally concentrating the CARDs. In NLRP1-CT, UPA dimerization or oligomerization is made apparent by an intrinsic propensity for CARD-CARD dimerization along an interface that does not participate in classical type I, II, or III helical interactions, which we resolve as additional molecules decorating the central NLRP1^CARD^ filament. The exact functional role of NLRP1 CARD-CARD dimerization remains to be fully addressed.

The role of the UPA subdomain in UPA-CARD inflammasomes is analogous to other oligomerization domains in inflammasome sensors. For example, in the activated NLRC4 inflammasome, NBDs and LRRs mediate the oligomerization into a disk-shaped signalling platform, clustering the attached CARDs towards the centre of the assembly. The clustered CARDs then template the assembly of ASC and/or caspase-1 to stimulate inflammasome signalling^5,6,33,44^. NLRP3, which possesses a similar domain architecture, might also use a similar mechanism, as mutations in predicted NBD and LRR contacts abolished its inflammasome activity^24^.

Other immune sensing pathways also employ CARD filament assembly strategies. In the RIG-I RNA sensing pathway, oligomerization of the RIG-I helicase domain on RNA clusters flexibly linked RIG-I 2CARDs, inducing their tetramerization. This RIG-I 2CARD tetramer, which represents one full helical turn, templates the assembly of the CARD filament of the MAVS adaptor to stimulate interferon signalling^36,45,46^. In the CARMA1/BCL10/MALT1 (CBM) signalosome, CARMA1^CARD^ nucleates the formation of a BCL10 filament through CARD-CARD interactions. However, the presence of an additional CARMA1 domain called the coiled coil region, which stimulates CARMA1 self-oligomerization, enhances BCL10 filament formation^37,47^. The role of the RIG-I helicase domain and the CARMA1 coiled coil region are thus similar to that of the UPA subdomain, by which self-oligomerization domains cluster flexibly linked CARDs to stimulate filament formation and signalling complex assembly.

Nucleated filament growth underlies the assembly of inflammasomes^3^. Such behaviour allows signal amplification from a low concentration of a pathogen- or danger-activated sensor that acts as a nucleator to a robust host response^42^. Our data are consistent with this model, whereby CARD8^CARD^ directly recruits and polymerizes caspase-1^CARD^ through the favourable side of the growing filament seed. NLRP1^CARD^, however, must recruit the adapter ASC to facilitate caspase-1 interaction and polymerization^30^. The specificity of the ASC filament for NLRP1^CARD^ on one side and caspase-1^CARD^ on the other makes it a perfect bridge between the NLRP1 sensor and the caspase-1 effector, and additional signalling amplification might occur through ASC^PYD^ polymerization. Thus, the ability to oligomerize and the specificity of the hetero-oligomerization determine the composition of the particular pathways and their biological outcomes.

## Methods

### Cloning, protein expression and purification

For structure determination, the UPA-CARD fragments of CARD8 (S297-L537, T60 isoform, Uniprot ID: Q9Y2G2-5) and NLRP1 (S1213-S1473, isoform 1, Uniprot ID: Q9C000-1) were cloned into a modified pDB-His-MBP vector with an HRV 3C protease linker (His-MBP-3C-UPA-CARD). For *in vitro* filament formation experiments, the CARDs of CARD8 (P446-L537) and NLRP1 (D1374-S1473) were also cloned into the modified pDB-His-MBP vector. For *in vitro* analyses, expression constructs encoding CARD8^CARD^ and NLRP1^CARD^ had an additional A452C (α1) or G1463C (α6) mutation, respectively, for cysteine labelling applications. For cellular studies, the NLRP1 CT was cloned into pcDNA3.1 LIC 6A with a C-terminal FLAG tag (UPA-CARD-FLAG) and the CARD8 CT was cloned pcDNA3.1 LIC 6B encoding a C-terminal mCherry tag (UPA-CARD-mCherry). Site-directed mutagenesis was performed with Q5 polymerase (NEB).

cDNA encoding ASC^CARD^ (106-195) and caspase-1^CARD^ (1-95) connected by a 5XGSS linker was cloned into the pDB-His-MBP vector with a tobacco etch virus (TEV)-cleavable N-terminal His6-MBP tag using NdeI and Not I restriction sites. Mutations of W169G on ASC^CARD^ (Type IIa) and G20K on caspase-1^CARD^ (Type IIb) were performed by QuikChange Mutagenesis (Agilent Technologies). All plasmids were confirmed by Sanger sequencing. All constructs will be deposited on Addgene.

To generate NLRP1 CT filaments, constructs were transformed into BL21 (DE3) cells, grown to OD600 0.6-0.8, cold shocked on ice water for 20 min, and induced overnight with 0.4 mM Isopropyl β-D-1-thiogalactopyranoside (IPTG) at 18 °C. The cells were harvested by centrifugation (4000 RPM, 20 min) and lysed by sonication (2 s on 2 s off, 4 min total on, 55% power, Branson) in lysis buffer (25 mM Tris-HCl pH 7.5, 150 mM NaCl, 1 mM TCEP, SIGMAFAST protease inhibitor). Cell lysate was then centrifuged (17,000 RPM, 1 h) and the soluble proteome was incubated with pre-equilibrated amylose resin for 2 h at 4 °C. Bound resin was then washed by gravity flow with 50 column volume (CV) lysis buffer and eluted with 3 CV elution buffer (25 mM Tris-HCl pH 7.5, 150 mM NaCl, 1 mM TCEP, 50 mM maltose). Eluate from amylose resin was concentrated and purified as a range of small oligomers on a Superose 6 size exclusion column (Cytiva) in lysis buffer, concentrated again (500 μL, A_280_ = 6) using a 50 kDa spin column (Millipore), and cleaved overnight with MBP-3C protease (50 μL, A280 = 3) on ice. These cleavage conditions were optimized extensively to produce elongated filaments, as uncleaved MBP-tagged protein incorporated in the NLRP1-CT filaments limits polymerization. Filaments were further purified by size exclusion chromatography on a Superose 6 column (Cytiva) in lysis buffer and recovered from the column void fraction at a suitable concentration for cryo-EM (A_280_ = 0.4).

To express and purify CARD8 CT filaments, expression and purification protocol followed as above until after elution from maltose resin. Eluate was concentrated (500 μL, A_280_ = 6) using a 50 kDa spin column (Millipore). Following extensive optimization, the eluate was cleaved with MBP-3C protease (50 μL, A_280_ = 3) for 4 h at 37 °C to produce elongated filaments. Filaments were further purified by size exclusion chromatography on a Superose 6 column (Cytiva) in lysis buffer, recovered from the column void fraction, and slowly concentrated using a 0.5 mL spin concentrator (Millipore, 100 kDa MW cutoff).

To obtain monomeric MBP-CARD proteins, constructs were expressed and purified similarly to the UPA-CARD filaments. Following elution from amylose resin, monomeric MBP-CARD proteins were concentrated to 1 mL (Amicon Ultra, 50 kDa MW cutoff) and further purified by size exclusion chromatography on a Superdex 200 column (Cytiva) in lysis buffer.

Expression of ASC^CARD^-CASP1^CARD^ was similar to above. Cells were then collected and resuspended in lysis buffer (25 mM Tris-HCl at pH 8.0, 150 mM NaCl, 20 mM imidazole 5 mM β-ME), followed by sonication. The cell lysate was clarified by centrifugation at 40,000 *g* at 4 °C for 30 min. The clarified supernatant containing the target protein was incubated with pre-equilibrated Ni-NTA resin (Qiagen) for 30 min at 4 °C. After incubation, the resin–supernatant mixture was poured into a gravity column and the resin was washed with lysis buffer. The protein was eluted using lysis buffer supplemented with 500 mM imidazole. The Ni-NTA eluate was then incubated with TEV protease at 16 °C overnight, and the cleaved His6-MBP tag was removed by passing the protein through an amylose resin column (Qiagen). The flow-through fraction containing the tag-free protein was further purified using a Superdex 200 (10/300 GL) gel-filtration column (Cytiva). Purified ASC^CARD^-CASP1^CARD^ oligomer was assessed for absolute molecular mass by re-injecting the sample into the Superdex 200 column equilibrated with 25 mM Tris-HCl at pH 8.0, 150 mM NaCl, and 2 mM DTT. The chromatography system was coupled to a three-angle light scattering detector (mini-DAWN TRISTAR) and a refractive index detector (Optilab DSP) (Wyatt Technology). Data were collected every 0.5 s with a flow rate of 0.5 mL/min. Data analysis was carried out using ASTRA V.

### Negative stain EM

Copper grids coated with layers of plastic and thin carbon film (Electron Microscopy Sciences) were glow discharged before 5 μl of purified proteins (A_280_ = 0.1) were applied. Samples were left on the grids for 1 min, blotted, and then stained with 1% uranyl formate for 30 s, blotted, and air dried. The grids were imaged on a JEOL 1200EX or Tecnai G^2^ Spirit BioTWIN microscope at the Harvard Medical School (HMS) EM facility operating at 80 keV.

### Cryo-EM data collection of NLRP1 and CARD8 filaments

Cryo-EM data collection of NLRP1 CT filaments was conducted at the HMS Cryo-EM Center. Purified NLRP1 CT filaments (A_280_ = 0.50; 25 mM Tris-HCl pH 7.5, 150 mM NaCl, 1 mM TCEP) were loaded onto a glow-discharged C-flat grid (CF-1.2/1.3 400-mesh copper-supported holey carbon, Electron Microscopy Sciences), blotted for 4–5 s under 100% humidity at 4 °C, and plunged into liquid ethane using a Mark IV Vitrobot (ThermoFisher). Grids were screened for ice and particle quality prior to data collection. 1,988 movies were acquired using a Titan Krios microscope (ThermoFisher) at an acceleration voltage of 300 keV equipped with a BioQuantum K3 Imaging Filter (slit width 25 eV), and a K3 direct electron detector (Gatan) operating in counting mode at 81,000 x (1.06 Å pixel size). Automated data collection with SerialEM varied the defocus range between −0.8 to −2.2 μm with four holes collected per stage movement through image shift. All movies were exposed with a total dose of 52.3 e^−^/Å^2^ for 3.5 s fractionated over 50 frames.

Cryo-EM data collection of CARD8 CT filaments was conducted at the Pacific Northwest Center for Cryo-EM (PNCC). Purified CARD8 CT filaments (A_280_ = 0.75; 25 mM Tris-HCl pH 7.5, 150 mM NaCl, 1 mM TCEP) were loaded onto a glow-discharged Quantifoil copper grid (R1.2/1.3 400-mesh copper-supported holey carbon, Electron Microscopy Sciences), blotted for 4–5 s under 100% humidity at 4 °C, and plunged into liquid ethane using a Mark IV Vitrobot (ThermoFisher). Grids were screened for ice and particle quality prior to data collection. 1,208 movies were acquired using a Titan Krios microscope (ThermoFisher) at an acceleration voltage of 300 keV equipped with a Falcon 3EC direct electron detector (ThermoFisher) operating in counting mode at 96,000 x (0.8315 Å pixel size). Automated data collection with EPU varied the defocus range between −0.8 to −2.2 μm with three movies collected per hole through image shift prior to stage movement. All movies had an exposure time of approximately 40 s for a total dose of 40 e^−^/Å^2^ fractionated over 50 frames.

The CARD octamer sample was applied to a glow discharged quantifoil holey carbon R1.2/1.3 copper grid and plunged with a Mark IV Vitrobot (ThermoFisher) set to 4°C and 100% humidity. Cryo-EM data were collected at the UMass cryo-EM facility on a Talos Arctica microscope with a K2 Summit® direct electron detection camera. 1,606 movies were collected in super-resolution mode, with an accumulated dose of ~58 e^−^/Å^2^ fractionated into 50 frames. The physical pixel size was 1.17 Å, and the defocus range was 1.2-2.6 μm.

### Cryo-EM data processing of NLRP1 and CARD8 filaments

For the CARD8 CT filament, 1,208 movies were aligned using RELION’s implementation of the MotionCor2 algorithm^48^. The defocus values and contrast transfer functions (CTFs) of the motion-corrected micrographs were then computed and corrected for using CTTFFIND4.1^49^. 1,181 micrographs were selected for further processing based on a maximum resolution criterion of 10 Å. Subsequently, start-end particle coordinates were manually picked and 287,568 particles were extracted in RELION^50^, with a box size of 640 pixels, shift of 6 Å for each segment box, and 2 times binning^50,51^ (320 pixel box). After 2D classification, 163,233 particles remained. An initial helical symmetry of −99° and 5.2 Å was derived from the power spectrum of the best 2D class average. An initial model was built with relion_helix_inimodel2d^52^ using this 2D class and the initial helical parameters as input. 111,333 particles were selected after 3D classification, which were re-extracted without binning and used for final 3D refinement. The refined helical symmetry was −99.07° and 5.20 Å. Postprocessing in RELION led to a 3.54 Å reconstruction (Figure 1, Supplementary Fig. 1).

For the NLRP1 CT filament, movie frames were aligned using RELION’s implementation of the MotionCor2 algorithm^48^. The defocus values and CTFs of the motion-corrected micrographs were then computed and corrected for using CTTFFIND4.1^49^. 1,578 micrographs were selected for further processing based on a maximum CTF resolution criterion of 5 Å. Subsequently, start-end particle coordinates were manually picked, and 118,006 particles were extracted in RELION^50^ with a box size of 512 pixels and shift of 16 Å for each segment box^50,51^. All particles were downscaled to a box size 360 to reduce processing time. After 2D classification, 55,083 particles remained. An initial helical symmetry of −100.8° and 5.07 Å was derived from the power spectrum of the best 2D class average. An initial model (helical lattice) was built with relion_helix_toolbox^52^ using these initial helical parameters as input. 18,272 particles were selected after 3D classification and used for final 3D refinement. The refined helical symmetry was −100.79° and 5.08 Å. Postprocessing in RELION led to a 3.72 Å reconstruction (Figure 1, Supplementary Fig. 2).

For the ASC^CARD^-caspase-1^CARD^ octamer, motion correction of the raw movies was performed with MotionCor2 ^48^ and the averaged micrographs were binned 3 times over the super-resolution pixel size, resulting a pixel size of 1.75 Å. The CTF parameters were calculated with Gctf^53^. 351,275 particles were picked with samautopick.py, and subjected to 2D classification in Relion^50^. 303,336 particles remained after 2D classification. After initial 3D refinement, a homemade script was used to remove redundant particles from dominant views, and 89,861 particles remained. Per particle CTF parameters were refined with Gctf prior to the final refinement. Postprocessing in RELION led to a 5.0 Å reconstruction (Supplementary Fig. 5).

### Model building and display

Model building was performed in program Coot^54^. The monomeric NLRP1^CARD^ or CARD8^CARD^ crystal structure (PDB ID: 4IFP^39^ or 4IKM^40^) was fit to the NLRC4^CARD^ filament model (PDB ID: 6N1I^33^) and used as initial models for refinement. Refinement was performed using Phenix^55^ and Refmac^56^. For the ASC^CARD^-caspase-1^CARD^ octamer, subunit structures from ASC^CARD^ (PDB ID: 6N1H^33^) and caspase-1^CARD^ filaments (PDB ID: 5FNA^7^) were fitted individually into the octamer cryo-EM map, and refined using Refmac^56^. Structural representations were displayed and rendered using Pymol^57^, UCSF ChimeraX^58^, and UCSF Chimera^59^. Session files will be made available on our Open Science Framework (https://osf.io/x7dv8/), and raw data will be made available on EMPIAR.

### Cell death assay

HEK 293T cells stably expressing CASP1 and GSDMD-V5 were seeded at 2 x 10^5^ cells/well in 12-well tissue culture dishes. The following day, cells were transfected with plasmids encoding for the indicated NLRP1 or CARD8 construct (0.02 μg), ASC for NLRP1 experiments (0.01 μg), and RFP (to 1 μg) using FuGENE HD according to manufacturer’s instructions (Promega). 24 h later supernatants were analysed for LDH activity using the Pierce LDH Cytotoxicity Assay Kit (ThermoFisher) and lysates were evaluated by immunoblotting for caspase-1 (Cell Signaling Technology, #2225), GSDMD (Abcam, Ab9116) and GAPDH (Cell Signaling Technology, 14C10).

### Confocal Imaging

HEK 293T cultured in Dulbecco’s modified Eagle’s medium (DMEM) supplemented with 10% fetal bovine serum (FBS) were plated on CELLview 4-compartment dishes (Greiner Bio-One). HEK 293T cells were transfected with CARD8 WT and mutant UPA-CARD-mCherry constructs (0.02 μg) using lipofectamine 2000 (Thermo Fisher Scientific). 48 hours after transfection, cells were fixed with 4% PFA for 10 minutes at RT. Cells were imaged using a spinning disk confocal Nikon microscope equipped with a Plan Apo 20x/1.3 air objective. Image analysis was performed in Fiji^60^.

### Data availability

The cryo-EM structures have been deposited in the Electron Microscopy Data Bank (EMDB) with accession numbers EMD-22219 (CARD8-CT filament), (NLRP1-CT filament), and EMDB-22233 (ASC^CARD^-caspase-1^CARD^ octamer). The atomic coordinates have been deposited in the protein databank (PDB) with accession numbers 6XKJ (CARD8-CT filament) and 6XKK (NLRP1-CT filament). Pymol/chimera session files will be made available on our Open Science Framework (https://osf.io/x7dv8/), and raw data will be made available on EMPIAR.

## Acknowledgements

We thank Maria Ericsson at the HMS EM facility for advice and training, Richard Walsh, Sarah Sterling, and Shaun Rawson at the Harvard Cryo-EM Center for Structural Biology for cryo-EM training and productive consultation, Paula Montero Llopis at the Microscopy Resources on the North Quad (MicRoN) core at Harvard Medical School for microscope use, Theo Humphreys for cryo-EM screening and data collection at the Pacific Northwest Center for Cryo-EM (PNCC), Chen Xu and Kangkang Song for cryo-EM screening and data collection at UMass Medical School, BioRender for figure design, and SBGrid for computing support. This work was supported by National Institutes of Health grants DP1 HD087988 and R01 AI124491 to H.W., T32 GM007739-Andersen and F30 CA243444 to A.R.G., the Memorial Sloan Kettering Cancer Center Core Grant P30 CA008748 to D.A.B.. A portion of this research was also supported by NIH grant U24GM129547 and performed at the PNCC at OHSU and accessed through EMSL (grid.436923.9), a DOE Office of Science User Facility sponsored by the Office of Biological and Environmental Research.

## Author contributions

H.W., L.R.H., and H.S. conceived the project idea and designed the study. L.R.H. designed constructs. L.R.H., H.S., P.F., and T.M.F. carried out preliminary expression and purification studies. L.R.H., P.F., and L.D. purified the filaments. T.M. F., L.R.H., L.D., and P.F. made cryo-EM grids for data collection. L.R.H. screened grids and collected cryo-EM data. Y.L., L.D., and L.R.H. analysed cryo-EM data, and L.R.H., and L.D. performed model building and refinement. L.R.H. and L.D. designed and cloned mutants for in vitro and cell-based assays of the filaments. L.D. performed in vitro filament formation experiments and confocal microscopy. A.R.G. performed cellular signalling experiments under the supervision of D.A.B. J.R. designed, expressed and purified the ASC-caspase-1 octamer and collected its cryo-EM data. L.R.H. and H.W. wrote the manuscript with input from all authors.

## Additional information

**Supplementary Information** See below

---

**Supplementary Figure 1.**
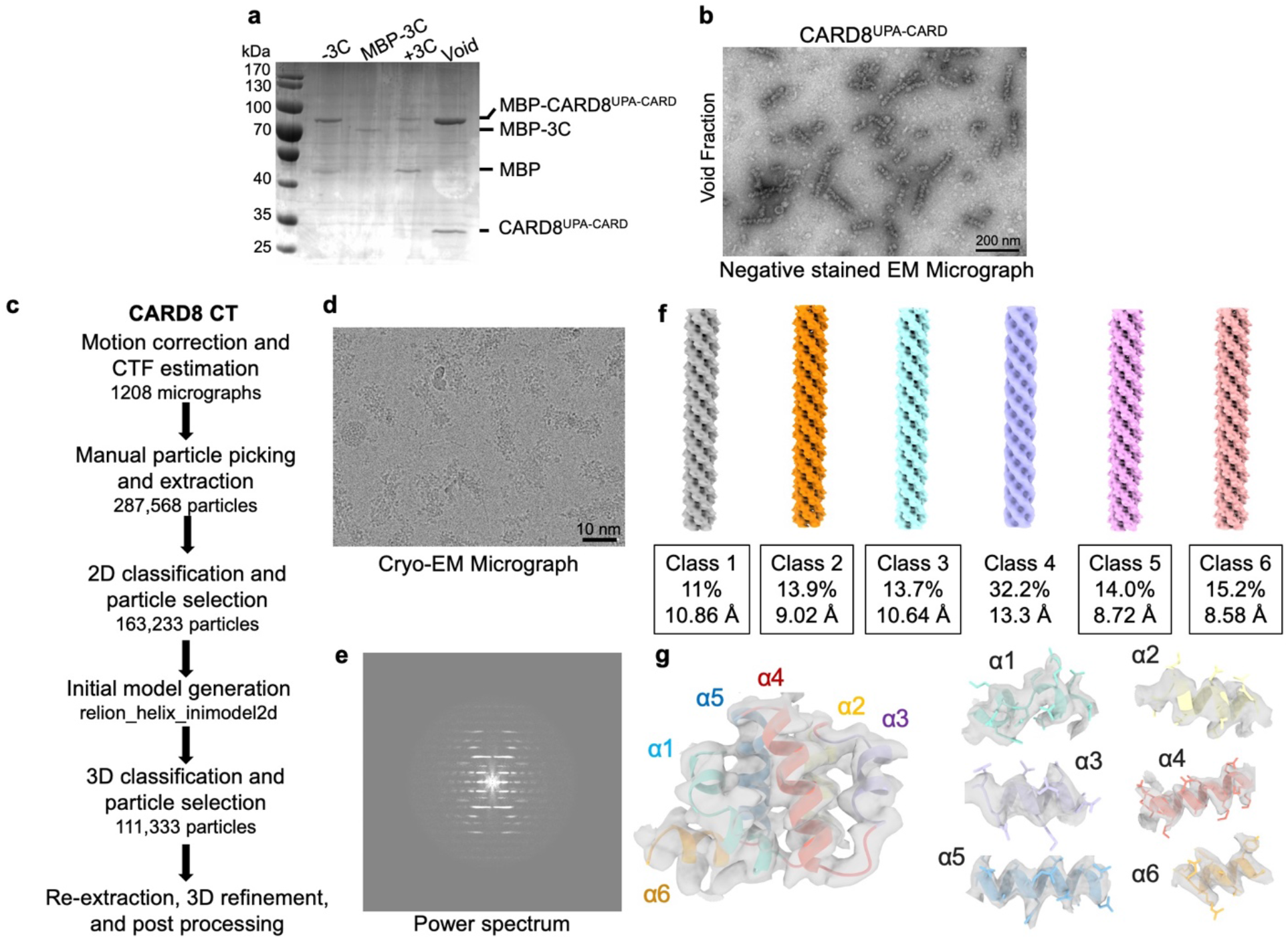
Structure determination of the CARD8-CT filament. (**a**) SDS-PAGE showing purification of the CARD8^UPA-CARD^ oligomer. Some uncleaved protein incorporates into the growing filaments and is present in the column void fraction. (**b**) Negative stain EM of the column void fraction showing short CARD-CT filaments. (**c**) Data processing flow chart for structure determination in RELION. (**d**) Representative cryo-EM micrograph of CARD8-CT filaments. (**e**) Power spectrum derived from a single 2D class average. (**f**) 3D classes generated during data processing. The chosen classes for further refinement are indicated by a box. (**g**) Fit of the atomic model of the CARD8^CARD^ core filament into the cryo-EM density.

**Supplementary Figure 2.**
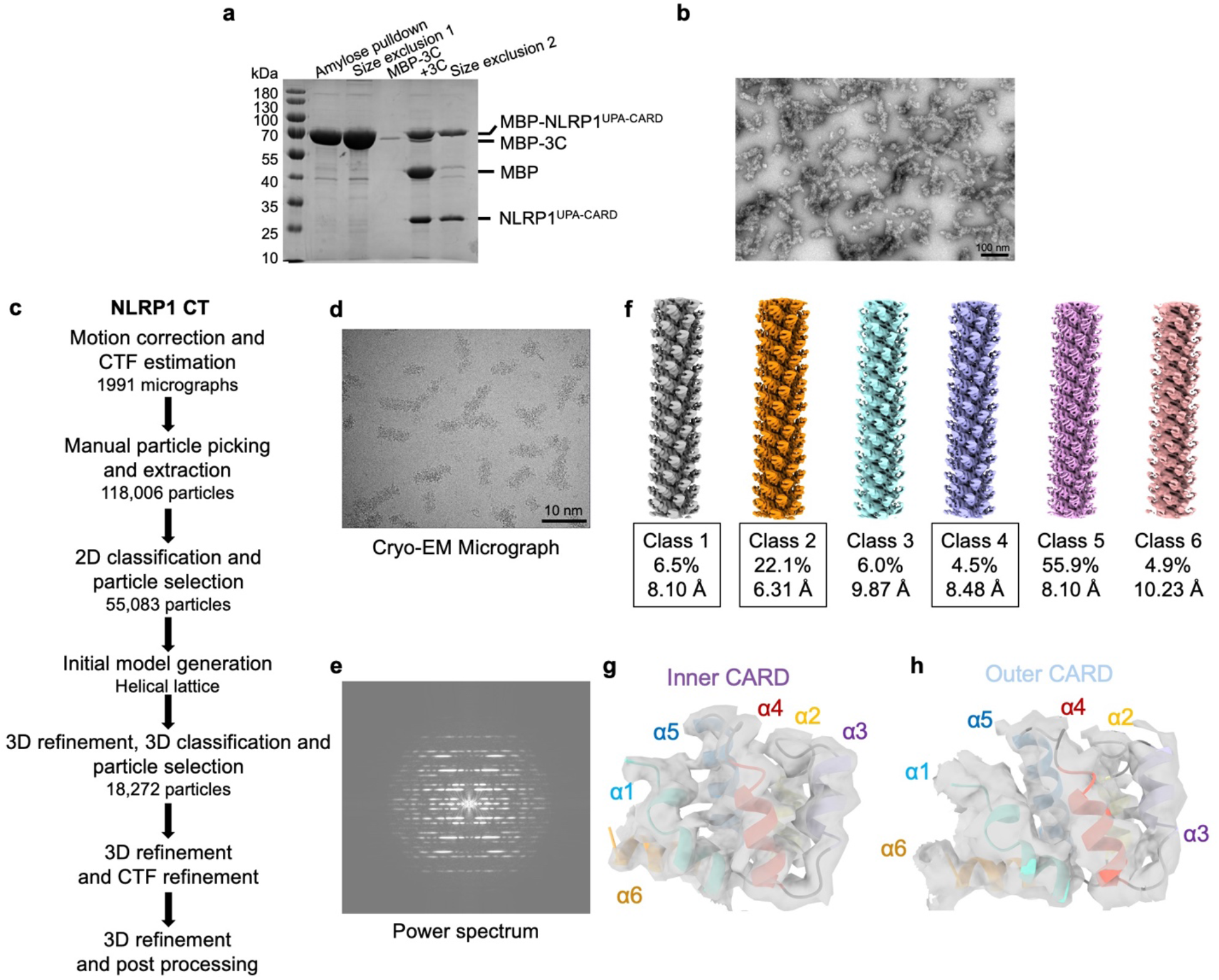
Structure determination of the NLRP1-CT filament. (**a**) SDS-PAGE showing purification of the NLRP1^UPA-CARD^ oligomer. Some uncleaved protein incorporates into the growing filaments and is present in the column void fraction. (**b**) Negative stain EM of the column void fraction showing short NLRP1-CT filaments. (**c**) Data processing flow chart for structure determination in RELION. (**d**) Representative cryo-EM micrograph of the NLRP1-CT filaments. (**e**) Power spectrum derived from a single 2D class average. (**f**) 3D classes generated during data processing. The chosen classes for further refinement are indicated by a box. (**g-h**) Fit of the atomic model of the inner NLRP1^CARD^ (**g**) and the outer NLRP1^CARD^ (h) in the cryo-EM density.

**Supplementary Figure 3.**
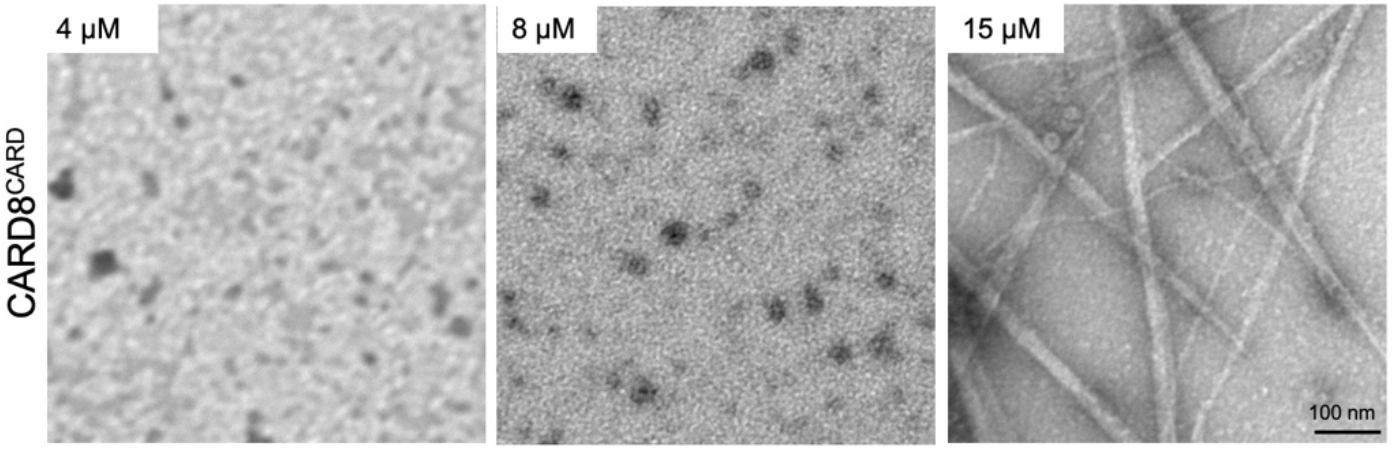
Determination of the critical concentration for CARD8^CARD^ filament formation using negative staining EM. Purified monomeric MBP-3C-CARD8^CARD^ was cleaved at concentrations ranging from 1-15 μM. CARD8^CARD^ filaments did not appear consistently until 15 μM. Once formed, CARD8^CARD^ filaments had the tendency to bundle.

**Supplementary Figure 4.**
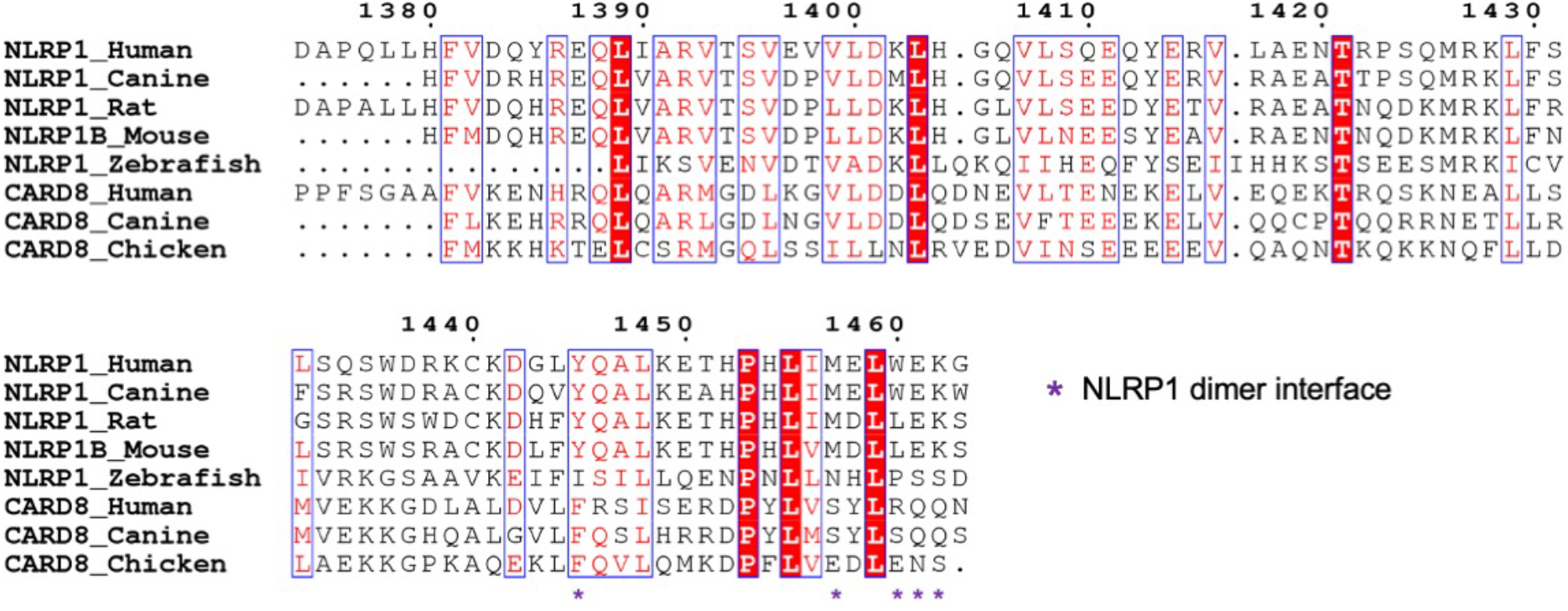
Sequence conservation of the NLRP1 dimer interface. Multiple sequence alignment (Clustal Omega) between NLRP1^CARD^ and CARD8^CARD^ homologs, coloured with ESPrint^62^. While the NLRP1 dimer interface (annotated with purple asterisks) is conserved across NLRP1 paralogs, these residues differ in CARD8.

**Supplementary Figure 5.**
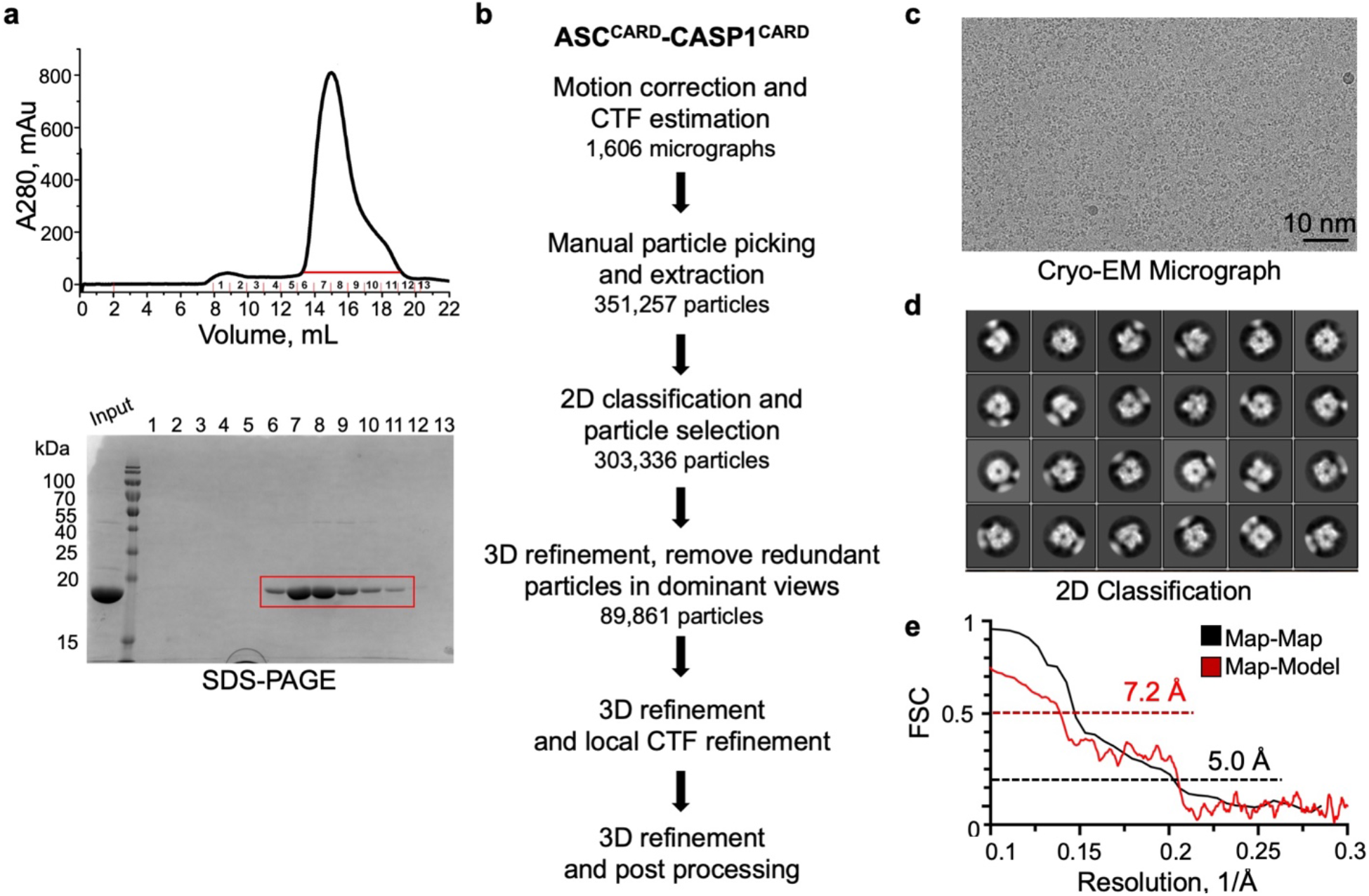
Structure determination of the ASC-caspase-1 octamer. (**a**) Purification of the ASC-caspase-1 octamer. The indicated fractions were run on an SDS-PAGE gel. (**b**) Data processing flow chart for structure determination in RELION. (**c**) Representative cryo-EM micrograph. (**d**) Representative 2D class averages. (**e**) Gold-standard FSC and map-model correlation plots of the ASC-caspase-1 octamer, which gave an overall resolution of 5.0 Å.

**Supplementary Table 1.**
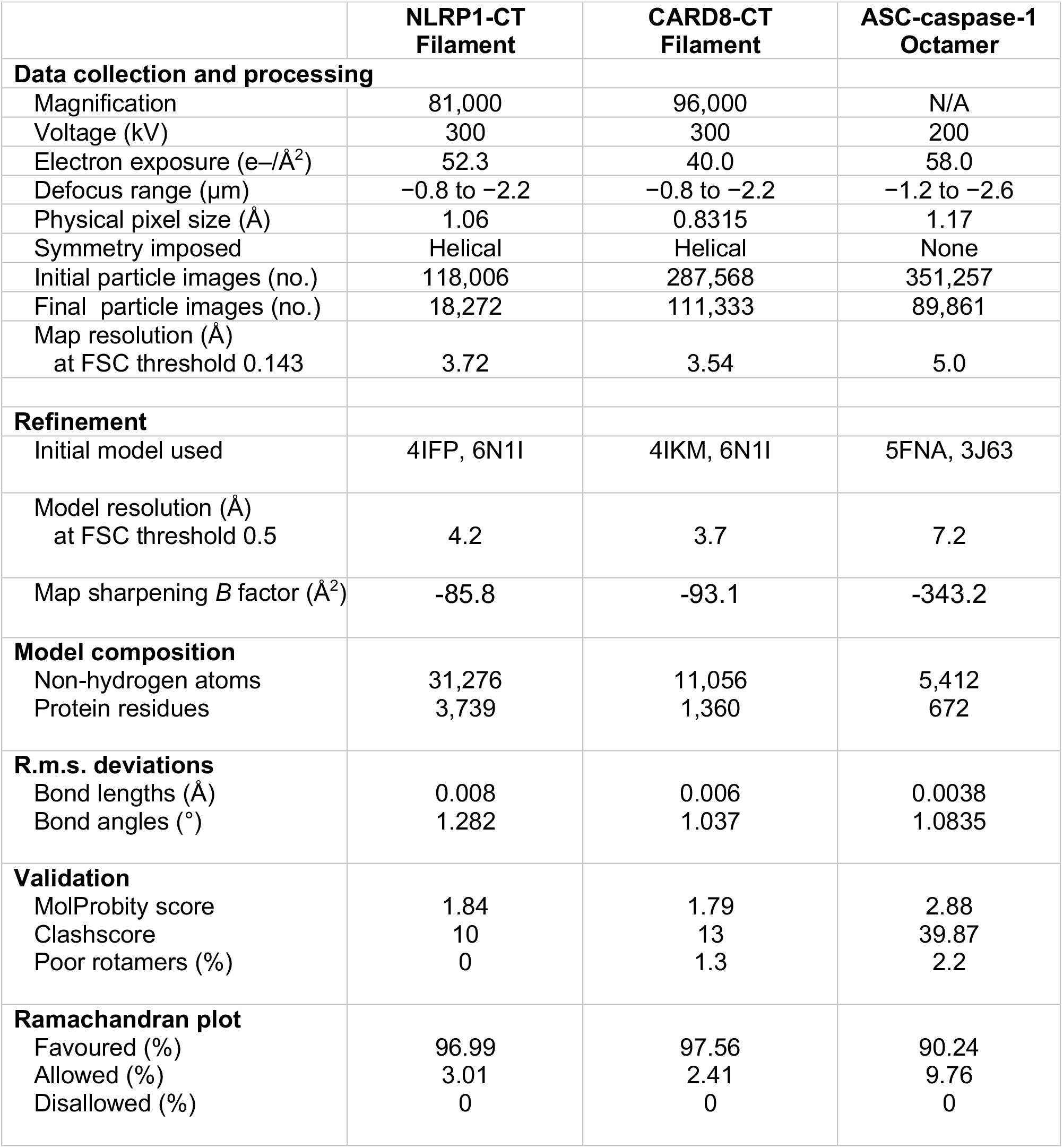
Cryo-EM data collection and refinement statistics.

## References

1 Martinon, F., Burns, K. & Tschopp, J. The Inflammasome: A Molecular Platform Triggering Activation of Inflammatory Caspases and Processing of proIL-β. Molecular Cell 10, 417–426, (2002).

2 Broz, P. & Dixit, V. M. Inflammasomes: mechanism of assembly, regulation and signalling. Nat Rev Immunol 16, 407–420, (2016).

3 Lu, A. et al. Unified Polymerization Mechanism for the Assembly of ASC-Dependent Inflammasomes. Cell 156, 1193–1206, (2014).

4 Shen, C., Sharif, H., Xia, S. & Wu, H. Structural and mechanistic elucidation of inflammasome signaling by cryo-EM. Curr Opin Struct Biol 58, 18–25, (2019).

5 Hu, Z. et al. Structural and biochemical basis for induced self-propagation of NLRC4. Science 350, 399–404, (2015).

6 Zhang, L. et al. Cryo-EM structure of the activated NAIP2-NLRC4 inflammasome reveals nucleated polymerization. Science 350, 404–409, (2015).

7 Lu, A. et al. Molecular basis of caspase-1 polymerization and its inhibition by a new capping mechanism. Nat Struct Mol Biol 23, 416–425, (2016).

8 Park, H. H. et al. The death domain superfamily in intracellular signaling of apoptosis and inflammation. Annu Rev Immunol 25, 561–586, (2007).

9 Kesavardhana, S., Malireddi, R. K. S. & Kanneganti, T.-D. Caspases in Cell Death, Inflammation, and Pyroptosis. Annual Review of Immunology 38, 567–595, (2020).

10 Lieberman, J., Wu, H. & Kagan, J. C. Gasdermin D activity in inflammation and host defense. Sci Immunol 4, (2019).

11 Xia, S., Hollingsworth, L. R. & Wu, H. in Cell Survival and Cell Death (eds K. Newton, J. M. Murphy, & E. A. Miao) (CSHL Press, Cold Spring Harb Perspect Biol, 2019).

12 Ruan, J., Xia, S., Liu, X., Lieberman, J. & Wu, H. Cryo-EM structure of the gasdermin A3 membrane pore. Nature 557, 62–67, (2018).

13 Taabazuing, C. Y., Griswold, A. R. & Bachovchin, D. A. The NLRP1 and CARD8 inflammasomes. Immunological Reviews n/a, (2020).

14 Mitchell, P. S., Sandstrom, A. & Vance, R. E. The NLRP1 inflammasome: new mechanistic insights and unresolved mysteries. Current Opinion in Immunology 60, 37–45, (2019).

15 Finger, J. N. et al. Autolytic Proteolysis within the Function to Find Domain (FIIND) Is Required for NLRP1 Inflammasome Activity. Journal of Biological Chemistry 287, 25030–25037, (2012).

16 D’Osualdo, A. et al. CARD8 and NLRP1 Undergo Autoproteolytic Processing through a ZU5-Like Domain. PLOS ONE 6, e27396, (2011).

17 Zhong, F. L. et al. Germline NLRP1 Mutations Cause Skin Inflammatory and Cancer Susceptibility Syndromes via Inflammasome Activation. Cell 167, 187–202.e117, (2016).

18 Xu, H. et al. The N-end rule ubiquitin ligase UBR2 mediates NLRP1B inflammasome activation by anthrax lethal toxin. The EMBO Journal, e101996, (2019).

19 Chui, A. J. et al. N-terminal degradation activates the NLRP1B inflammasome. Science 364, 82–85, (2019).

20 Sandstrom, A. et al. Functional degradation: A mechanism of NLRP1 inflammasome activation by diverse pathogen enzymes. Science 364, eaau1330, (2019).

21 Zhong, F. L. et al. Human DPP9 represses NLRP1 inflammasome and protects against auto-inflammatory diseases via both peptidase activity and FIIND domain binding. Journal of Biological Chemistry 293, 18864–18878, (2018).

22 Okondo, M. C. et al. Inhibition of Dpp8/9 Activates the Nlrp1b Inflammasome. Cell chemical biology 25, 262–267.e265, (2018).

23 Johnson, D. C. et al. DPP8/DPP9 inhibitor-induced pyroptosis for treatment of acute myeloid leukemia. Nature medicine 24, 1151–1156, (2018).

24 Sharif, H. et al. Structural mechanism for NEK7-licensed activation of NLRP3 inflammasome. Nature, (2019).

25 Levandowski, C. B. et al. NLRP1 haplotypes associated with vitiligo and autoimmunity increase interleukin-1β processing via the NLRP1 inflammasome. Proceedings of the National Academy of Sciences of the United States of America 110, 2952–2956, (2013).

26 Jin, Y. et al. NALP1 in vitiligo-associated multiple autoimmune disease. N Engl J Med 356, 1216–1225, (2007).

27 Grandemange, S. et al. A new autoinflammatory and autoimmune syndrome associated with NLRP1 mutations: NAIAD (NLRP1-associated autoinflammation with arthritis and dyskeratosis). Annals of the rheumatic diseases 76, 1191–1198, (2017).

28 Drutman, S. B. et al. Homozygous NLRP1 gain-of-function mutation in siblings with a syndromic form of recurrent respiratory papillomatosis. Proc Natl Acad Sci U S A 116, 19055–19063, (2019).

29 Srinivasula, S. M. et al. The PYRIN-CARD Protein ASC Is an Activating Adaptor for Caspase-1. Journal of Biological Chemistry 277, 21119–21122, (2002).

30 Ball, D. P. et al. Caspase-1 interdomain linker cleavage is required for pyroptosis. Life Sci Alliance 3, (2020).

31 Poyet, J. L. et al. Identification of Ipaf, a human caspase-1-activating protein related to Apaf-1. J Biol Chem 276, 28309–28313, (2001).

32 Broz, P. et al. Redundant roles for inflammasome receptors NLRP3 and NLRC4 in host defense against Salmonella. J Exp Med 207, 1745–1755, (2010).

33 Li, Y. et al. Cryo-EM structures of ASC and NLRC4 CARD filaments reveal a unified mechanism of nucleation and activation of caspase-1. Proceedings of the National Academy of Sciences 115, 10845–10852, (2018).

34 Matyszewski, M. et al. Cryo-EM structure of the NLRC4CARD filament provides insights into how symmetric and asymmetric supramolecular structures drive inflammasome assembly. Journal of Biological Chemistry, (2018).

35 Gong, X. et al. Structural basis for distinct inflammasome complex assembly by human NLRP1 and CARD8. bioRxiv doi: https://doi.org/10.1101/2020.06.17.156307, (2020).

36 Wu, B. et al. Molecular imprinting as a signal-activation mechanism of the viral RNA sensor RIG-I. Mol Cell 55, 511–523, (2014).

37 David, L. et al. Assembly mechanism of the CARMA1-BCL10-MALT1-TRAF6 signalosome. Proc Natl Acad Sci U S A 115, 1499–1504, (2018).

38 Ferrao, R. & Wu, H. Helical assembly in the death domain (DD) superfamily. Curr Opin Struct Biol 22, 241–247, (2012).

39 Jin, T., Curry, J., Smith, P., Jiang, J. & Xiao, T. S. Structure of the NLRP1 caspase recruitment domain suggests potential mechanisms for its association with procaspase-1. Proteins 81, 1266–1270, (2013).

40 Jin, T., Huang, M., Smith, P., Jiang, J. & Xiao, T. S. The structure of the CARD8 caspase-recruitment domain suggests its association with the FIIND domain and procaspases through adjacent surfaces. Acta crystallographica. Section F, Structural biology and crystallization communications 69, 482–487, (2013).

41 Dick, M. S., Sborgi, L., Ruhl, S., Hiller, S. & Broz, P. ASC filament formation serves as a signal amplification mechanism for inflammasomes. Nat Commun 7, 11929, (2016).

42 Wu, H. Higher-Order Assemblies in a New Paradigm of Signal Transduction. Cell 153, 287–292, (2013).

43 Kagan, J. C., Magupalli, V. G. & Wu, H. Supramolecular organizing centres: location-specific higher-order signalling complexes that control innate immunity. Nat Rev Immunol 14, 821–826, (2014).

44 Diebolder, C. A., Halff, E. F., Koster, A. J., Huizinga, E. G. & Koning, R. I. Cryoelectron Tomography of the NAIP5/NLRC4 Inflammasome: Implications for NLR Activation. Structure (London, England : 1993) 23, 2349–2357, (2015).

45 Peisley, A., Wu, B., Yao, H., Walz, T. & Hur, S. RIG-I Forms Signaling-Competent Filaments in an ATP-Dependent, Ubiquitin-Independent Manner. Molecular Cell 51, 573–583, (2013).

46 Peisley, A., Wu, B., Xu, H., Chen, Z. J. & Hur, S. Structural basis for ubiquitin-mediated antiviral signal activation by RIG-I. Nature 509, 110–114, (2014).

47 Qiao, Q. et al. Structural Architecture of the CARMA1/Bcl10/MALT1 Signalosome: Nucleation-Induced Filamentous Assembly. Mol Cell 51, 766–779, (2013).

48 Zheng, S. Q. et al. MotionCor2: anisotropic correction of beam-induced motion for improved cryo-electron microscopy. Nat Methods 14, 331–332, (2017).

49 Mindell, J. A. & Grigorieff, N. Accurate determination of local defocus and specimen tilt in electron microscopy. Journal of Structural Biology 142, 334–347, (2003).

50 He, S. & Scheres, S. H. W. Helical reconstruction in RELION. Journal of Structural Biology 198, 163–176, (2017).

51 Zivanov, J. et al. New tools for automated high-resolution cryo-EM structure determination in RELION-3. eLife 7, e42166, (2018).

52 Scheres, S. Amyloid structure determination in RELION-3.1. Acta Crystallographica Section D 76, 94–101, (2020).

53 Zhang, K. Gctf: Real-time CTF determination and correction. Journal of Structural Biology 193, 1–12, (2016).

54 Emsley, P., Lohkamp, B., Scott, W. G. & Cowtan, K. Features and development of Coot. Acta crystallographica. Section D, Biological crystallography 66, 486–501, (2010).

55 Adams, P. D. et al. PHENIX: a comprehensive Python-based system for macromolecular structure solution. Acta Crystallographica Section D 66, 213–221, (2010).

56 Murshudov, G. N. et al. REFMAC5 for the refinement of macromolecular crystal structures. Acta crystallographica. Section D, Biological crystallography 67, 355–367, (2011).

57 Schrodinger, LLC. The PyMOL Molecular Graphics System, Version 1.8 (2015).

58 Goddard, T. D. et al. UCSF ChimeraX: Meeting modern challenges in visualization and analysis. Protein Science 27, 14–25, (2018).

59 Pettersen, E. F. et al. UCSF Chimera—A visualization system for exploratory research and analysis. J. Comput. Chem. 25, 1605–1612, (2004).

60 Schindelin, J. et al. Fiji: an open-source platform for biological-image analysis. Nat Methods 9, 676–682, (2012).

61 Kucukelbir, A., Sigworth, F. J. & Tagare, H. D. Quantifying the local resolution of cryo-EM density maps. Nat Methods 11, 63–65, (2014).

62 Robert, X. & Gouet, P. Deciphering key features in protein structures with the new ENDscript server. Nucleic acids research 42, W320–W324, (2014).

